# Incomplete immunity in a natural animal-microbiota interaction boosts pathogen virulence

**DOI:** 10.1101/2023.09.20.558495

**Authors:** Kim L. Hoang, Timothy D. Read, Kayla C. King

**Affiliations:** Department of Biology, University of Oxford; Oxford, UK; Emory University School of Medicine, Division of Infectious Diseases; Atlanta, Georgia, USA; Department of Zoology, University of British Columbia; Vancouver, British Columbia, Canada; Department of Microbiology & Immunology, University of British Columbia; Vancouver, British Columbia, Canada

## Abstract

Strong partial immunity in recovered hosts is predicted to favour more virulent pathogens upon re-infection in the population. We present empirical evidence that the incomplete immunity generated by commensal host microbiota can similarly select for higher pathogen virulence. We tracked the evolutionary trajectories of a widespread pathogen (*Pseudomonas aeruginosa*) experimentally passaged through populations of nematode hosts which had been immune-primed by a natural commensal. Immune protection selected for pathogens more than twice as likely to kill the nematode as those evolved in non-primed or immune-compromised animals. Despite the higher virulence that emerged, pathogen molecular evolution in immune-primed hosts was slower and more constrained compared to evolution in immune-compromised hosts, where substantial genetic differentiation was exhibited. These findings directly attribute the partial protective immunity provided by host-microbiome interactions as a significant selective force shaping the virulence and evolutionary dynamics of novel infectious diseases.

---

When an animal clears an infection, immune memory—a phenomenon that occurs in invertebrates and vertebrates—can protect against future infection^1^. Incomplete immunity occurs when a pathogen can re-infect, although the outcome is likely to result in reduced disease severity and death^2^. The commensal microbes colonising hosts (*i*.*e*., microbiota) can also generate a protective and long-lasting host immune response, even if the microbes themselves are cleared^3–5^. Heightened expression of defence genes in the host can be primed through detection of microbe-associated molecular patterns found in both pathogens and microbiota^6^. This is a common mechanism in nature by which host microbiota can help against infectious disease^7–9^. While direct interactions between commensal microbes and pathogens can select for lower virulence^10,11^, immune-mediated mechanisms may have the opposite effect if pathogen colonisation can still occur^4,12,13^. Incomplete immunity can reduce the costs of virulence to pathogens, an outcome which suggests the leakiness of infection-induced immune protection might favour more virulent pathogens^14^. It is unclear whether incomplete immunity from host-microbiota interactions can similarly drive the evolution of pathogens which cause higher host mortality.

## Results and Discussion

To directly test whether host microbiota can shape pathogen virulence via immune responses, we experimentally evolved a widespread, disease-causing animal pathogen (*Pseudomonas aeruginosa*) upon introduction to a natural host-commensal interaction. *Caenorhabditis elegans* nematodes can be infected by the bacterium *P. aeruginosa*, which harms them by accumulating in the host intestine and destroying tissue over time^15^. Although nematodes are found naturally with *Pseudomonas* spp.^16^, the isolate used here (PA14) was from burn wounds in humans^17^ and thus novel to *C. elegans*. These nematodes are however frequently associated with a commensal species, *P. berkeleyensis*^*18,19*^. Although this microbiota member is ultimately cleared upon pathogen exposure, its initial colonisation induces genes regulated by a mitogen-activated protein kinase (MAPK)—an ancient innate immune pathway found in plants and animals^18,20^—thereby enhancing nematode host survival during *P. aeruginosa* infection (^18^ and Fig 1A). By comparison, immune-compromised mutants (a knock-out of the MAPK ortholog) were killed readily by *P. aeruginosa*, with no protective effect elicited by *P. berkeleyensis* colonisation (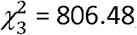, P < 0.001, Fig. 1A). The immunity conferred by *P. berkeleyensis* for wild-type hosts was incomplete as the pathogen can form a stable infection in protected hosts but had a lower load (Student’s t = 7.02, P < 0.001, Fig. 1B). *Pseudomonas berkeleyensis* is mildly pathogenic in the absence of threat, similar to other protective microbes (F_(3,8)_ = 11.28, P = 0.003; Figure S1)^21,22^. Thus, consistent with earlier work using vaccines^23^ and vertebrate-infectious disease interactions^14^, nematode immunity here reduced the costs of virulence to *P. aeruginosa* by protecting hosts from the disease-induced mortality that would likely limit onward transmission^24^.

**Fig. 1.**
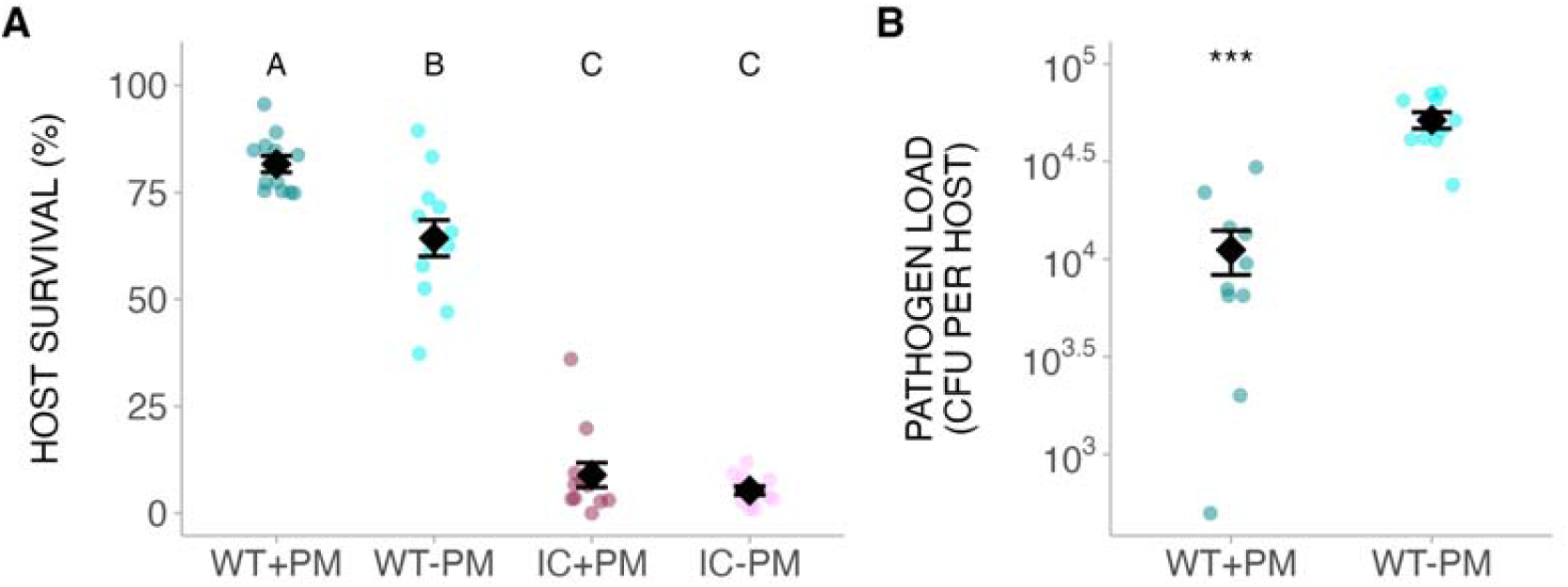
Host microbiota provides incomplete immune protection. **(A)** Host survival (mean ± SE) upon pathogen infection with or without exposure to microbiota member. **(B)**. Pathogen load (mean ± SE) in each host. WT = wild-type host, IC = immunocompromised host, PM = protective microbiota (*P. berkeleyensis*). Different letters indicate significant differences. ***P < 0.001

We experimentally passaged pathogen populations independently in nematode populations either previously colonized by *P. berkeleyensis* or in naïve (non-primed) populations (Fig. 2A). These treatments were conducted alongside a no-host control for lab adaptation, as well as in nematode mutants (*pmk-1*) not capable of mounting the primed immune response (Fig. 2A). We carried out phenotypic assays of host mortality upon infection (metric for pathogen virulence) and load (metric for pathogen fitness) across pathogen generations and treatments. We then used shotgun sequencing of pools of 40 colonies to measure evolutionary changes in the genomic composition of *P. aeruginosa* populations.

**Fig. 2.**
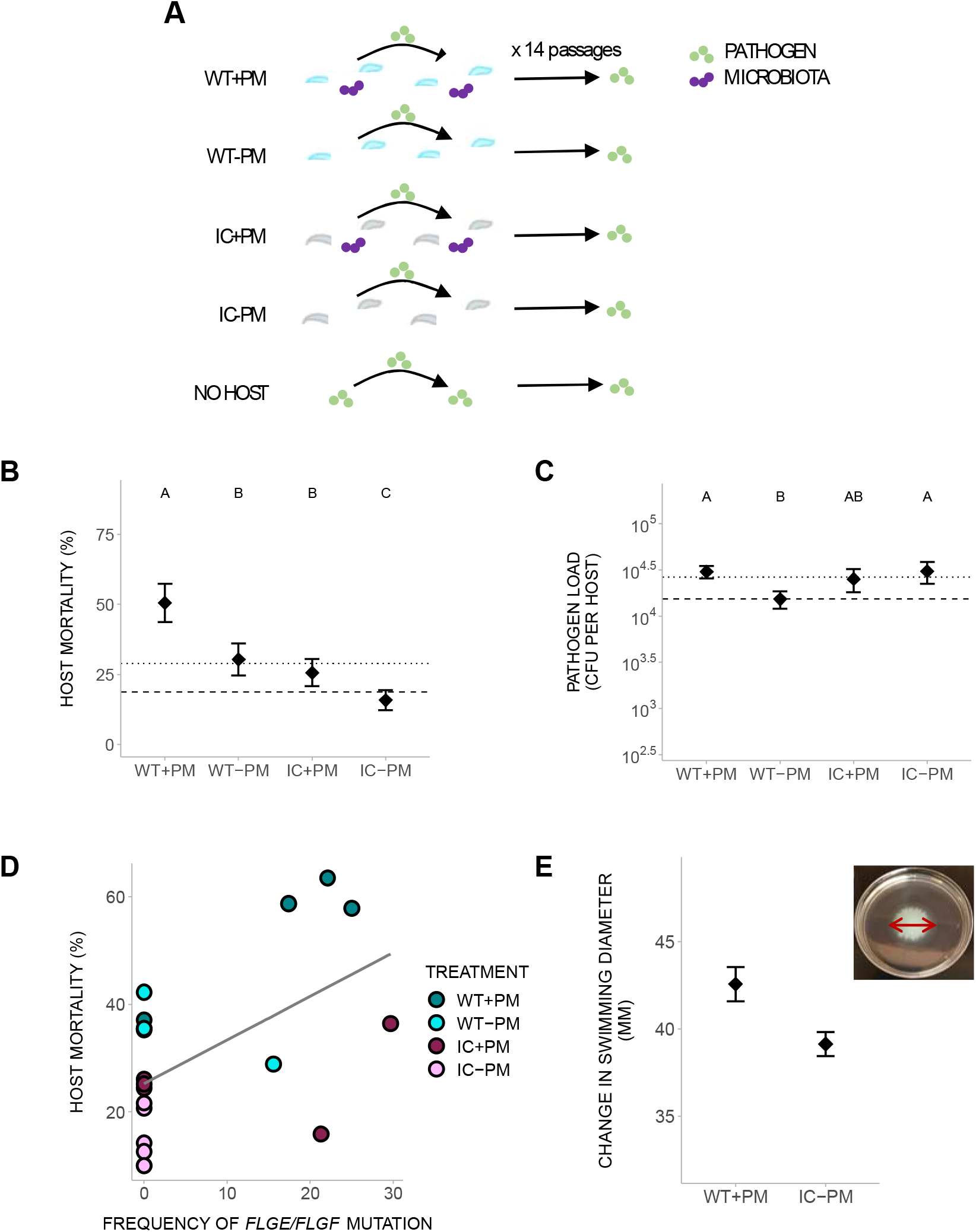
Incomplete immunity from microbiota selects for more virulent pathogens. **(A)** Experimental evolution design. WT = wild-type host, IC = immunocompromised host, PM = protective microbiota (*P. berkeleyensis*, purple dots), green dots = pathogen (P. aeruginosa). **(B)** Mortality of wild-type hosts without microbiota when infected with evolved pathogens (x-axis). **(C)** Load of evolved pathogens (x-axis) in wild-type hosts without microbiota. Shaded dashed line indicates mean ± SE for hosts infected by no-host control pathogen. Dotted line indicates mean for hosts infected by ancestral pathogen. **(D)** Correlation between host mortality and frequency of a flagellar-related (*flgE/flgF*) mutation. **(E)** Swimming motility of most virulent and least virulent pathogens. (inset) Example of bacterial diameter measured for swimming motility assessment. All error bars are mean ± SE. Different letters indicate significant differences.

Microbiota-induced incomplete immunity selected for more virulent pathogens compared to naïve hosts (= 55.39, P < 0.001, Fig. 2B). These findings support theoretical models on incomplete immunity generated from prior pathogen exposure and vaccines^14,23^. That microbiota in an invertebrate host can affect pathogens similarly to antibody-generating vaccines and pathogen exposure in vertebrate hosts points to a more general role of incomplete immunity in virulence evolution, regardless of the specific priming mechanism. Despite the boost in virulence, pathogens had not evolved to overcome the protective effects of immune priming (microbiota: = 2.36, P = 0.12; host: = 0.066, P = 0.80; interaction: = 2.35, P = 0.13; Fig. S2A). Immune priming can still offer harm-reduction from increasingly virulent pathogens able to colonise (Fig. S2B). Hosts exposed to *P. berkeleyensis* selected for reduced virulence in naïve immune-compromised hosts (= 6.31, P = 0.01; Fig. S3), but there were no significant host or interaction effects (host: = 0.08, P = 0.78; interaction: = 2.55, P = 0.11). This result points to a trade-off in virulence for pathogens evolving in primed hosts. These pathogens had the highest virulence in naïve immunocompetent hosts relative to other evolved pathogens, but lower virulence in naïve immune-compromised hosts. Collectively, our phenotypic findings demonstrate that the immediate benefits of increased survival and pathogen tolerance conferred by the microbiota can ultimately lead to extremely negative impacts on the host^25^.

Pathogen virulence and load evolved along different trajectories. The levels of host mortality caused during infection and bacterial accumulation per host were not correlated across treatments (Fig. 2C). This result corroborates previous research showing virulence in novel pathogens can evolve along independent trajectories in experimental replicates^26,27^. We hypothesized that density-independent virulence factors, such as toxin production or motility, may be contributors to the higher virulence emerging in pathogens from immune-primed hosts. To identify potential targets of selection on virulence mechanisms, we pool-sequenced evolved pathogen populations (see Methods) and quantified the mutations arising from the ancestor. Each population had 400-500 mutations, with most partially increasing to <50% of the population (Figs. S4A and S4B). Several mutations had fixed in the same genes or intergenic regions across treatments, with several genes having multiple mutations (Figs. S5 and S6). Further pairwise comparisons between treatments revealed allele frequency differences in genes involved in diverse biological pathways (Figs. S7 and S8), and treatment replicates had few unique mutations in common (Fig. S9). These results suggest that virulence under selection in our experiment has a polygenic basis, as found in other pathogens with broad host ranges^28–30^.

We compared the population genomic composition between treatments with the largest difference in evolved virulence (i.e., immune-primed vs. naïve, immune-compromised hosts, Fig. 2B). We found an intergenic mutation between two genes involved in bacterial flagella function (*flgE/flgF*). Flagellar proteins are notable inducers of immune responses, and alterations in regulatory regions are less likely to disrupt function^31,32^. Mutation frequency across replicates was positively correlated with infected host mortality (Spearman’s rho = 0.49, P = 0.03; Figs. 2D and S10). Since flagella are virulence factors^33,34^ and are necessary for motility, we compared the swimming ability of evolved populations (see Methods and Fig. 2 inset). Pathogen motility significantly differed between these extreme treatments (Fig. 2E), although differences across all treatments were marginally insignificant (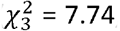, P = 0.052, Fig. S10B). Whilst flagellar efficiency may be partially contributing to high pathogen virulence, disruption in metabolism may also be playing a role in the reduced virulence^33^ evolved in immune-compromised hosts. A mutation prominent across treatments and negatively correlated with host mortality (Figs. S10C and S10D) was in the *fmt* (methionyl-tRNA formyltransferase) gene responsible for translation initiation^35^. Our results suggest that while *P. aeruginosa* utilized different genetic pathways to adapt to immune-primed and immune-compromised hosts, both groups converged on similar fitness levels^36,37^.

The strength of the host immune response induced by microbiota can shape genomic evolution in novel pathogens. Pathogen replication in the presence of weak selection—such as exhibited in immune-compromised hosts—can make it easier for mutations to accumulate, resulting in extensive genomic diversification from the ancestor. Such rapid changes in genome evolution has been shown in bacterial pathogens responsible for zoonotic diseases^38,39^ as well as viral pathogens^40^. The initial lower pathogen load in immune-primed hosts (Fig. 1B) may also dampen the number of new mutations that can be acquired in these populations^41^. We constructed phylogenies based on point mutations to assess the relationship between individual pathogen colonies and the ancestor (Fig. 3). Most mutations identified in each colony had fixed in the pooled samples (Fig. S11). Pathogens evolving in immune-compromised hosts diverged substantially from the ancestor (5.57 ± 0.80 mutations per colony; Figs. 3, S12, and S13). These colonies also shared similar distances from the ancestor as those evolving *in vitro*, in addition to converging on similar virulence levels (Figs. 2B and 3A). These results indicate that mutations acquired from weak selection can reduce virulence and increase genetic diversity as previously found in pathogens infecting hosts with defects in their immune system^38,42,43^, where less virulent pathogens may be able to better compete against more virulent ones^44^. In contrast, pathogens evolving in immune-primed hosts had maintained only moderate genetic distance from the ancestor (3.21 ± 0.46 mutations per colony; Figs. 3, S12, and S14), suggesting the phenotypes we observed were due to interactions of large effect mutations. Despite selecting strongly for high virulence, immune protection ultimately limited pathogen evolution at the molecular level.

**Fig. 3.**
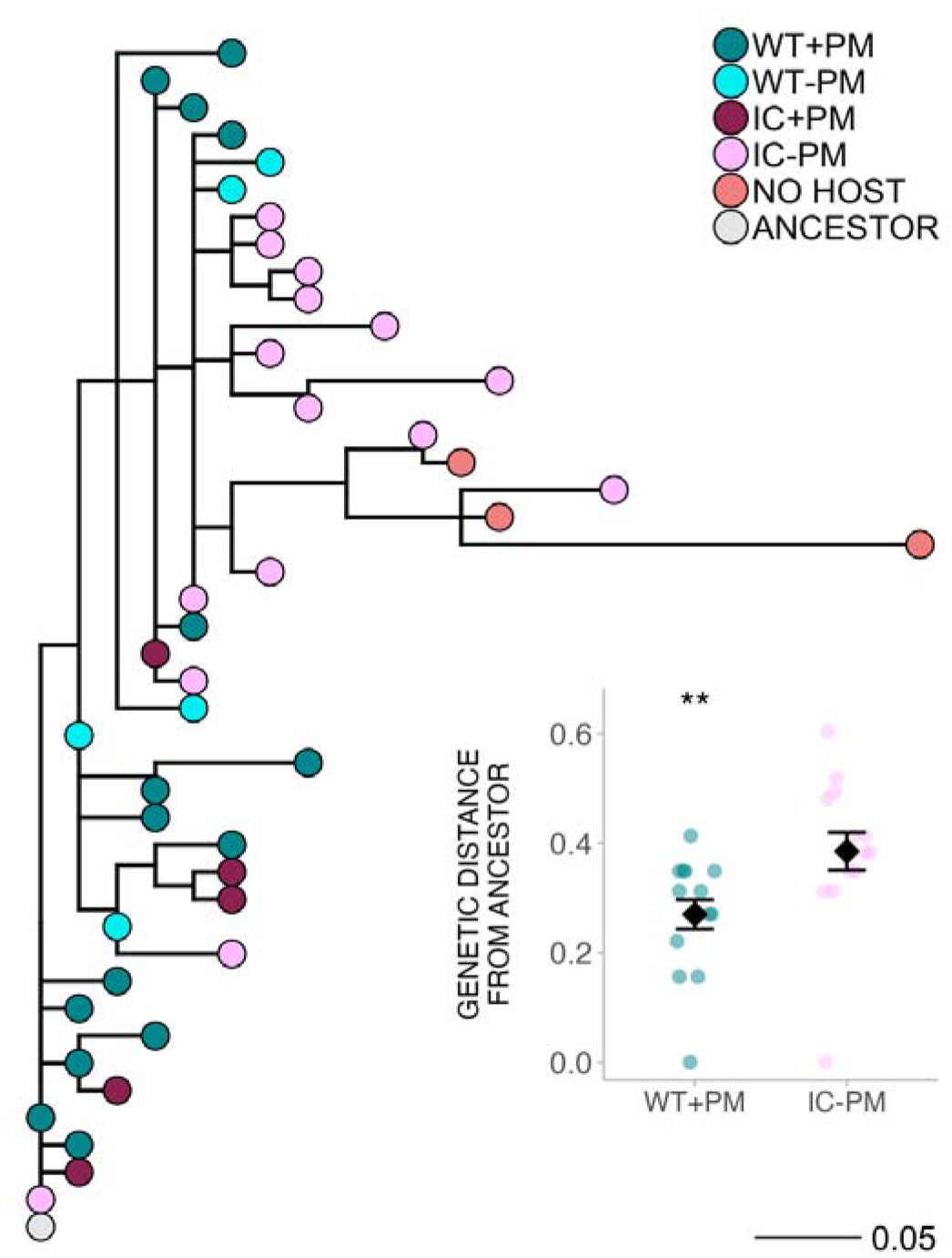
Incomplete immunity from microbiota constrains pathogen molecular evolution. Maximum parsimony phylogeny of colonies sampled from evolved pathogen populations. We sampled more colonies from the two treatments with the most contrast in virulence level: pathogens evolved in immune-primed hosts and naïve immune-compromised hosts. (inset) Genetic distance from the ancestor (mean ± SE) for colonies isolated from immune-primed hosts and naïve immune-compromised hosts. Values were square-root transformed to meet the condition for normal distribution. WT = wild-type host, IC = immunocompromised host, PM = protective microbiota. **P > 0.01

Immune responses can act to alter the degree of divergence between pathogen populations. Compromised host defences (i.e., weaker selection) may cause greater pathogen genetic divergence between populations compared to hosts with stronger defences^45–48^. We calculated pairwise F_ST_ for each SNP between replicate populations within each treatment to determine how host defence impacted pathogen population divergence (Fig. S15A). Pathogens evolving in immune-primed hosts had fewer significant F_ST_ loci compared to those evolving in hosts protected only by genome-encoded defence (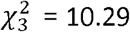, P = 0.016; Figs 4A and S15B). All treatments exhibited differentiation in genes involved in bacterial secretion system and two-component systems (Fig. S16). These results indicate that incomplete immune priming from host microbiota limited the genetic differentiation across replicate populations compared to in non-primed treatments.

**Fig. 4.**
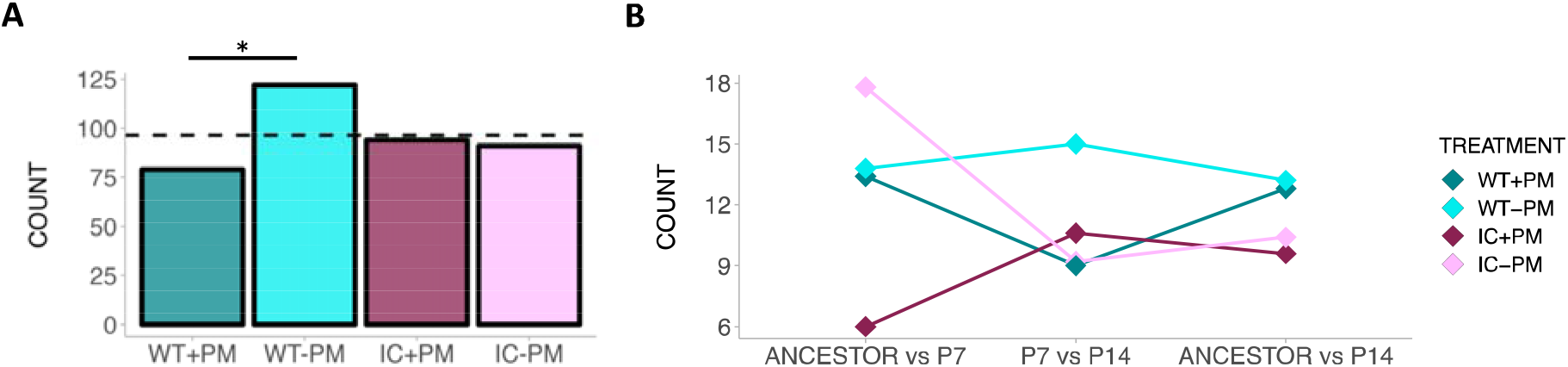
Host defences induced by microbiota alter pathogen evolutionary paths. (A and B) Count of loci between replicate populations at passage 14 (A) and between time points (B) within each treatment with significant genetic differentiation (FST). Dashed line indicates theoretical expectation. WT = wild-type host, IC = immunocompromised host, PM = protective microbiota. P7 = passage 7, P14 = passage 14. *P > 0.05

We also evaluated temporal shifts in the genetic composition of the whole population by calculating F_ST_ between the ancestral pathogen and evolved populations at the midpoint (i.e., passage seven) and endpoint of the experiment. At the midpoint, pathogens evolving in naïve hosts had more significant F_ST_ loci compared to those evolving in immune-primed hosts (F_3,16_ = 5.34, P = 0.010; Figure 4B; Fig. S17A). By the end of the experiment, treatments no longer varied in terms of the number of significant F_ST_ loci (F_3,16_ = 0.77, P = 0.53; Figs. 4B, S17B, and S17C). Earlier in evolutionary time, the absence of commensal microbiota generated more genetic differences between the ancestor and evolved pathogens, but eventually all populations exhibited similar rates of change. Taken together, the results suggest that the dynamics shaping pathogen evolution at the very beginning of emergence can become different after a period of adaptation^39,41^.

Host microbiota have been found to play significant roles in protecting hosts from harmful infection across the tree of life^9,49,50^. Over evolutionary time, however, we found that the incomplete immune protection induced by host microbiota can act similarly to vaccines^23,51^ and previous infection^14^ in favouring highly virulent pathogens. Conversely, immune-compromised hosts may serve as environments where pathogens can accumulate mutations, leading to genome degradation and host-restriction^38^. Host microbiota-immune interactions might therefore be a major source of selection shaping the ongoing evolution of emerging infectious diseases. Usage of probiotic microbes is becoming more prevalent across agricultural and wild systems^52,53^, including those species at risk of extinction due to rapid pathogen spread^54,55^. For long-lived hosts, application of probiotic microbes is a powerful tool to combat infectious diseases^56,57^. Identifying the mechanisms by which these microbes protect seems crucial to predicting their longer-term sustainability ‘in the field’. The efficacy of these therapeutics may be preserved despite pathogen evolution. However, proper precautions should be taken before potentially facilitating the generation of more virulent pathogens, balancing future risks with the immediate benefits to host individuals.

## References

1. Milutinović, B., and Kurtz, J. (2016). Immune memory in invertebrates. Semin. Immunol. 28, 328–342.

2. Leon, A.E., and Hawley, D.M. (2017). Host Responses to Pathogen Priming in a Natural Songbird Host. Ecohealth 14, 793–804.

3. Kim, Y., and Mylonakis, E. (2012). Caenorhabditis elegans immune conditioning with the probiotic bacterium Lactobacillus acidophilus strain ncfm enhances gram-positive immune responses. Infect. Immun. 80, 2500–2508.

4. Kwong, W.K., Mancenido, A.L., and Moran, N.A. (2017). Immune system stimulation by the native gut microbiota of honey bees. R. Soc. Open Sci. 4, 170003.

5. Clarke, T.B., Davis, K.M., Lysenko, E.S., Zhou, A.Y., Yu, Y., and Weiser, J.N. (2010). Recognition of peptidoglycan from the microbiota by Nod1 enhances systemic innate immunity. Nat. Med. 16, 228–231.

6. Selosse, M.A., Bessis, A., and Pozo, M.J. (2014). Microbial priming of plant and animal immunity: Symbionts as developmental signals. Trends Microbiol. 22, 607–613.

7. Hoang, K.L., and King, K.C. (2022). Symbiont-mediated immune priming in animals through an evolutionary lens. Microbiology, 1–11.

8. Gabrieli, P., Caccia, S., Varotto-Boccazzi, I., Arnoldi, I., Barbieri, G., Comandatore, F., and Epis, S. (2021). Mosquito trilogy: microbiota, immunity and pathogens, and their implications for the control of disease transmission. Front. Microbiol. 12, 1–17.

9. Khosravi, A., and Mazmanian, S.K. (2013). Disruption of the gut microbiome as a risk factor for microbial infections. Curr. Opin. Microbiol. 16, 221–227.

10. Ford, S.A., Kao, D., Williams, D., and King, K.C. (2016). Microbe-mediated host defence drives the evolution of reduced pathogen virulence. Nat. Commun. 7, 1–9.

11. Nelson, P., and May, G. (2020). Defensive symbiosis and the evolution of virulence. Am. Nat. 196, 333–343.

12. Horak, R.D., Leonard, S.P., and Moran, N.A. (2020). Symbionts shape host innate immunity in honeybees. Proc. R. Soc. B Biol. Sci. 287.

13. Corby-Harris, V., Snyder, L., Meador, C.A.D., Naldo, R., Mott, B., and Anderson, K.E. (2016). Parasaccharibacter apium, gen. Nov., sp. Nov., Improves Honey Bee (Hymenoptera: Apidae) resistance to Nosema. J. Econ. Entomol. 109, 537–543.

14. Fleming-Davies, A.E., Williams, P.D., Dhondt, A.A., Dobson, A.P., Hochachka, W.M., Leon, A.E., Ley, D.H., Osnas, E.E., and Hawley, D.M. (2018). Incomplete host immunity favors the evolution of virulence in an emergent pathogen. Science (80-.). 359, 1030–1033.

15. Irazoqui, J.E., Troemel, E.R., Feinbaum, R.L., Luhachack, L.G., Cezairliyan, B.O., and Ausubel, F.M. (2010). Distinct pathogenesis and host responses during infection of C. elegans by P. aeruginosa and S. aureus. PLoS Pathog. 6, 1–24.

16. Dirksen, P., Marsh, S.A., Braker, I., Heitland, N., Wagner, S., Nakad, R., Mader, S., Petersen, C., Kowallik, V., Rosenstiel, P., et al. (2016). The native microbiome of the nematode Caenorhabditis elegans: gateway to a new host-microbiome model. BMC Biol., 1–16.

17. Grace, A., Sahu, R., Owen, D.R., and Dennis, V.A. (2022). Pseudomonas aeruginosa reference strains PAO1 and PA14: A genomic, phenotypic, and therapeutic review. Front. Microbiol. 13, 1–15.

18. Montalvo-Katz, S., Huang, H., Appel, M.D., Berg, M., and Shapira, M. (2013). Association with soil bacteria enhances p38-dependent infection resistance in Caenorhabditis elegans. Infect. Immun. 81, 514–520.

19. Dirksen, P., Assié, A., Zimmermann, J., Zhang, F., Tietje, A.M., Marsh, S.A., Félix, M.A., Shapira, M., Kaleta, C., Schulenburg, H., et al. (2020). CeMbio - The Caenorhabditis elegans microbiome resource. G3 Genes, Genomes, Genet. 10, 3025–3039.

20. Engelmann, I., and Pujol, N. (2010). Innate immunity in C. elegans. In Invertebrate Immunity, pp. 105–121.

21. Vorburger, C., Ganesanandamoorthy, P., and Kwiatkowski, M. (2013). Comparing constitutive and induced costs of symbiont-conferred resistance to parasitoids in aphids. Ecol. Evol. 3, 706–713.

22. Lukasik, P., Guo, H., Van Asch, M., Ferrari, J., and Godfray, H.C.J. (2013). Protection against a fungal pathogen conferred by the aphid facultative endosymbionts Rickettsia and Spiroplasma is expressed in multiple host genotypes and species and is not influenced by co-infection with another symbiont. J. Evol. Biol. 26, 2654–2661.

23. Read, A.F., Baigent, S.J., Powers, C., Kgosana, L.B., Blackwell, L., Smith, L.P., Kennedy, D.A., Walkden-Brown, S.W., and Nair, V.K. (2015). Imperfect vaccination can enhance the transmission of highly virulent pathogens. PLoS Biol. 13, 1–18.

24. Pike, V.L., Stevens, E.J., Griffin, A.S., and King, K.C. (2023). Within- and between-host dynamics of producer and non-producer pathogens. Parasitology, 1–33.

25. Smith, C.A., and Ashby, B. (2023). Tolerance-conferring defensive symbionts and the evolution of parasite virulence. Evol. Lett. 7, 262–272.

26. Tardy, L., Giraudeau, M., Hill, G.E., McGraw, K.J., and Bonneaud, C. (2019). Contrasting evolution of virulence and replication rate in an emerging bacterial pathogen. Proc. Natl. Acad. Sci. U. S. A. 116, 16927–16932.

27. Ekroth, A.K.E., Gerth, M., Stevens, E.J., Ford, S.A., and King, K.C. (2021). Host genotype and genetic diversity shape the evolution of a novel bacterial infection. ISME J. 15, 2146–2157.

28. Chen, H., Bright, R.A., Subbarao, K., Smith, C., Cox, N.J., Katz, J.M., and Matsuoka, Y. (2007). Polygenic virulence factors involved in pathogenesis of 1997 Hong Kong H5N1 influenza viruses in mice. Virus Res. 128, 159–163.

29. Le Clec’h, W., Chevalier, F.D., McDew-White, M., Menon, V., Arya, G.A., and Anderson, T.J.C. (2021). Genetic architecture of transmission stage production and virulence in schistosome parasites. Virulence 12, 1508–1526.

30. Caseys, C., Shi, G., Soltis, N., Gwinner, R., Corwin, J., Atwell, S., and Kliebenstein, D.J. (2021). Quantitative interactions: The disease outcome of Botrytis cinerea across the plant kingdom. G3 Genes, Genomes, Genet. 11.

31. Sharp, C., and Foster, K.R. (2022). Host control and the evolution of cooperation in host microbiomes. Nat. Commun. 13, 1–15.

32. Rossez, Y., Wolfson, E.B., Holmes, A., Gally, D.L., and Holden, N.J. (2015). Bacterial Flagella: Twist and Stick, or Dodge across the Kingdoms. PLoS Pathog. 11, 1–15.

33. Feinbaum, R.L., Urbach, J.M., Liberati, N.T., Djonovic, S., Adonizio, A., Carvunis, A.R., and Ausubel, F.M. (2012). Genome-wide identification of Pseudomonas aeruginosa virulence-related genes using a Caenorhabditis elegans infection model. PLoS Pathog. 8, 11.

34. Duan, Q., Zhou, M., Zhu, L., and Zhu, G. (2013). Flagella and bacterial pathogenicity. J. Basic Microbiol. 53, 1–8.

35. Guillon, J.M., Mechulam, Y., Schmitter, J.M., Blanquet, S., and Fayat, G. (1992). Disruption of the gene for Met-tRNA(f)/(Met) formyltransferase severely impairs growth of Escherichia coli. J. Bacteriol. 174, 4294–4301.

36. Bailey, S.F., Rodrigue, N., and Kassen, R. (2015). The effect of selection environment on the probability of parallel evolution. Mol. Biol. Evol. 32, 1436–1448.

37. Frickel, J., Feulner, P.G.D., Karakoc, E., and Becks, L. (2018). Population size changes and selection drive patterns of parallel evolution in a host-virus system. Nat. Commun. 9, 1–10.

38. Klemm, E.J., Gkrania-Klotsas, E., Hadfield, J., Forbester, J.L., Harris, S.R., Hale, C., Heath, J.N., Wileman, T., Clare, S., Kane, L., et al. (2016). Emergence of host-adapted Salmonella Enteritidis through rapid evolution in an immunocompromised host. Nat. Microbiol. 1, 1–6.

39. Launay, A., Wu, C.J., Dulanto Chiang, A., Youn, J.H., Khil, P.P., and Dekker, J.P. (2021). In vivo evolution of an emerging zoonotic bacterial pathogen in an immunocompromised human host. Nat. Commun. 12, 1–12.

40. Kemp, S.A., Collier, D.A., Datir, R.P., Ferreira, I.A.T.M., Gayed, S., Jahun, A., Hosmillo, M., Rees-Spear, C., Mlcochova, P., Lumb, I.U., et al. (2021). SARS-CoV-2 evolution during treatment of chronic infection. Nature 592, 277–282.

41. Day, T., Kennedy, D.A., Read, A.F., and Gandon, S. (2022). Pathogen evolution during vaccination campaigns. PLoS Biol. 20, e3001804.

42. Mongelli, V., Lequime, S., Kousathanas, A., Gausson, V., Blanc, H., Nigg, J., Quintana-Murci, L., Elena, S.F., and Saleh, M.C. (2022). Innate immune pathways act synergistically to constrain RNA virus evolution in Drosophila melanogaster. Nat. Ecol. Evol.

43. Jansen, G., Crummenerl, L.L., Gilbert, F., Mohr, T., Pfefferkorn, R., Thänert, R., Rosenstiel, P., and Schulenburg, H. (2015). Evolutionary transition from pathogenicity to commensalism: Global regulator mutations mediate fitness gains through virulence attenuation. Mol. Biol. Evol. 32, 2883–2896.

44. Råberg, L., De Roode, J.C., Bell, A.S., Stamou, P., Gray, D., and Read, A.F. (2006). The role of immune-mediated apparent competition in genetically diverse malaria infections. Am. Nat. 168, 41–53.

45. Cruickshank, T., and Wade, M.J. (2008). Microevolutionary support for a developmental hourglass: Gene expression patterns shape sequence variation and divergence in Drosophila. Evol. Dev. 10, 583–590.

46. Runemark, A., Brydegaard, M., and Svensson, E.I. (2014). Does relaxed predation drive phenotypic divergence among insular populations? J. Evol. Biol. 27, 1676–1690.

47. MacPherson, A., and Nuismer, S.L. (2017). The probability of parallel genetic evolution from standing genetic variation. J. Evol. Biol. 30, 326–337.

48. Scribner, M.R., Santos-Lopez, A., Marshall, C.W., Deitrick, C., Coopera, V.S., and Hogan, D.A. (2020). Parallel evolution of tobramycin resistance across species and environments. MBio 11.

49. King, K.C. (2019). Defensive symbionts. Curr. Biol. 29, R78–R80.

50. Kaltenpoth, M., and Engl, T. (2014). Defensive microbial symbionts in Hymenoptera. Funct. Ecol. 28, 315–327.

51. Barclay, V.C., Sim, D., Chan, B.H.K., Nell, L.A., Rabaa, M.A., Bell, A.S., Anders, R.F., and Read, A.F. (2012). The evolutionary consequences of blood-stage vaccination on the rodent malaria Plasmodium chabaudi. PLoS Biol. 10, 9.

52. Duar, R.M., Lin, X.B., Zheng, J., Martino, M.E., Grenier, T., Pérez-Muñoz, M.E., Leulier, F., Gänzle, M., and Walter, J. (2017). Lifestyles in transition: evolution and natural history of the genus Lactobacillus. FEMS Microbiol. Rev. 41, S27–S48.

53. McKenzie, V.J., Kueneman, J.G., and Harris, R.N. (2018). Probiotics as a tool for disease mitigation in wildlife: insights from food production and medicine. Ann. N. Y. Acad. Sci. 1429, 18–30.

54. Hoyt, J.R., Langwig, K.E., White, J.P., Kaarakka, H.M., Redell, J.A., Parise, K.L., Frick, W.F., Foster, J.T., and Kilpatrick, A.M. (2019). Field trial of a probiotic bacteria to protect bats from white-nose syndrome. Sci. Rep. 9, 1–9.

55. Bletz, M.C., Loudon, A.H., Becker, M.H., Bell, S.C., Woodhams, D.C., Minbiole, K.P.C., and Harris, R.N. (2013). Mitigating amphibian chytridiomycosis with bioaugmentation: Characteristics of effective probiotics and strategies for their selection and use. Ecol. Lett. 16, 807–820.

56. Cross, M.L. (2002). Microbes versus microbes: Immune signals generated by probiotic lactobacilli and their role in protection against microbial pathogens. FEMS Immunol. Med. Microbiol. 34, 245–253.

57. Ouwehand, A.C., Forssten, S., Hibberd, A.A., Lyra, A., and Stahl, B. (2016). Probiotic approach to prevent antibiotic resistance. Ann. Med. 48, 246–255.

