## Supplementary material for "Incomplete immunity in a natural animal-microbiota interaction boosts pathogen virulence": Methods and supplemental figures

**The file includes:**

Materials and Methods

Figs. S1 to S17

Materials and Methods

Nematode and bacterial strains

Wild-type *C. elegans* strain N2 Bristol and *E. coli* strain OP50 were provided by the Caenorhabditis Genetics Center, which is funded by the NIH Office of Research Infrastructure Programs (P40 OD010440). Immunocompromised *C. elegans* mutant *pmk-1* (M03F8.4(op497), pmk-1(km25)) was provided by Jonathan Hodgkin (University of Oxford). *Pseudomonas berkeleyensis* MSPm1 (previously identified as *P. mendocina*) was provided by Michael Shapira (University of California at Berkeley). Green fluorescently-labelled *P. aeruginosa* PA14 (PA14-GFP) was provided by Kevin Foster (University of Oxford).

Survival and CFU assays with ancestral *P. aeruginosa*

*Host survival*

Eggs from N2 and pmk-1 nematodes were collected, surface-sterilized, and age-synchronized following a standard sodium hypochlorite protocol (*58*). After hatching, about 200 L1 larvae were spotted onto either lawns of *P. berkeleyensis* or *E. coli* grown on NGM plates. These nematodes were incubated at 20˚C for two days. L4/young adults were then transferred to *P. aeruginosa* plates and kept at 20˚C*.* After three days, the number of live nematodes were determined by prodding nematodes with a platinum pick to determine signs of movement.

*Pathogen CFU*

Following the steps above to infect nematodes for three days, we followed a modified protocol from (*59*) to determine the pathogen load in infected nematodes. Briefly, ten nematodes per population were picked into and washed twice with cold M9 buffer containing 0.01% Triton X-100 (M9-T), then chilled on ice for ~30 minutes to stop peristalsis. We then added enough cold bleach such that the final concentration is 0.3% in the nematode/M9 mixture. After briefly mixing, the mixture was kept on ice for 10 minutes, then cold M9-T added to stop the bleaching process. Nematodes were washed once more with cold M9-T and supernatant plated to check for efficiency of bleaching. Under a dissecting scope, we pipetted 10 individuals into another tube containing zirconium beads in about 100ul M9-T. Samples were shaken in a bead beater for 2 minutes at 27 1/s in a TissueLyser. After brief centrifugation, serially diluted homogenates were spread onto LB plates and incubated. The number of colony forming units were quantified after two days.

*Host fecundity*

We followed the steps as above to rear N2 or pmk-1 nematodes on either *P. berkeleyensis* or *E. coli*. We quantified the number of offspring on the same day we measured host mortality for nematodes infected with *P. aeruginosa*.

*Statistics*

All statistical analyses below were carried out in R version 4.2.0.

Survival data were analysed using a generalized linear mixed mode with a binomial distribution followed by Tukey multiple-comparison tests to determine pairwise differences. Pathogen CFU data were analysed using a t-test, and fecundity data were analysed using an ANOVA.

Experimental evolution

We passaged *P. aeruginosa* PA14-GFP under five treatments (Figure 2A): four host treatments and one no host treatment. To start, one single clone of PA14-GFP was grown overnight in LB broth and spread onto nematode growth medium (*60*), with subsequent incubation at 30˚C for one day. About 1000 nematodes were transferred from their respective rearing plates (described below) onto the *P. aeruginosa* plates and incubated at 20˚C. Nematodes were washed off each plate after one day, rinsed three times with M9 buffer. Ten percent of the M9/nematode mixture were crushed using a BeadBeater, and homogenates were plated onto LB plates. After overnight incubation, we picked 100 colonies into broth to start the next passage. Each treatment consisted of five replicate rearing and *P. aeruginosa* plates across 14 passages.

Nematodes were kept evolutionarily static (*i.e.*, not evolving) throughout the experiment. N2 and pmk-1 populations were reared as described in the *Survival and CFU assays with ancestral P. aeruginosa host survival* section. L4/young adults were transferred to *P. aeruginosa* plates as described above. For each passage, eggs were collected from stock nematode populations that were regularly resurrected from -80˚C to limit accumulation of *de novo* mutations in host lineages throughout the experiment.

*Mortality and CFU assays with evolved P. aeruginosa*

Mortality and CFU assays for evolved populations follow similar protocols as those for ancestral *P. aeruginosa* (*Survival and CFU assays with ancestral P. aeruginosa* section). Assays were performed in triplicates. For Figures 2B and 2C, we infected N2 nematodes that had been reared on OP50. For Figure S2, we infected N2 nematodes reared on *P. berkeleyensis*. For Figure S3, we infected pmk-1 nematodes reared on *E. coli* and quantified mortality after two days instead of three days due to high mortality of these hosts. Mortality and CFU data were analysed using generalized linear mixed models (with a binomial distribution or Poisson distribution, respectively) followed by Tukey multiple-comparison tests to determine pairwise differences.

Swimming motility

To measure motility of ancestral and evolved *P. aeruginosa*, we followed the protocol from (*61*) to inoculate swimming motility plates. We incubated plates at 30˚C for one day as this was the temperature NGM plates were incubated before nematodes were put on the pathogen. We the diameter of bacterial growth on this day as the initial diameter. We then incubated plates at 20˚C for three days following the infection timeline for the host mortality assay, then measured the final diameter. The initial diameter was subtracted from the final diameter to obtain the change in swimming diameter. Data were analysed using a linear mixed model.

DNA extraction and sequencing

For pooled samples, we grew 40 individual colonies (*i.e.*, clones) for each replicate population separately overnight in LB broth, then standardized the OD_600_ of each clone before pooling them into one tube to perform DNA extraction. For single colony samples, we grew individual colonies separately in LB broth overnight, then performed DNA extraction. For pooled samples. We extracted genomic DNA using DNeasy Blood and Tissue Kit (Qiagen) following the manufacturer’s instructions. Sample libraries were prepared using the Illumina DNA Prep kit and sequenced on an Illumina NextSeq 2000. Sequence quality was assessed using FastQC (https://www.bioinformatics.babraham.ac.uk/projects/fastqc/) and sequences were trimmed using fastP (*62*). we sequenced all five replicate populations of each host treatment and three random populations from the no host treatment. For single colony samples, we sequenced three random clones from each population of the WT+PM and IC-PM treatments, and one random clone from each population of the other treatments.

The ancestral PA14-GFP clone was sequenced using Oxford Nanopore Technologies (ONT) in addition to Illumina for hybrid assembly. Quality control and adapter trimming was performed with bcl2fastq (*63*) and porechop (*64*) using default parameters for Illumina and ONT sequencing, respectively. Hybrid assembly with Illumina and ONT reads was performed with Unicycler (*65*), and the resulting assembly was annotated using the Bakta annotation pipeline (*66*). Coverage of mapped reads was calculated using Samtools (*67*). Each pooled sample had at least 200X coverage. Each single colony sample had at least 60X coverage.

Analysis of pooled samples

We called variants with the ancestor as the reference using the Breseq pipeline polymorphism mode with default parameters (*68*). We tested whether the frequencies of *flgE/flgF* and *fmt* mutations were correlated with mortality using Spearman’s rank correlation.

To determine the allele frequency differences between treatments, we pooled together the reads across all replicate populations for each treatment. We then used the Popoolation2 pipeline (*69*) to calculate the exact allele frequency differences and estimated significance using Fisher's Exact Test. For significant loci found in coding regions, we used the Database for Annotation, Visualization and Integrated Discovery (DAVID) (*70*) tool to map each gene to the Kyoto Encyclopedia of Genes and Genomes (KEGG) pathways (*71*).

To calculate the per SNP F_ST_ within each treatment, we used the Popoolation2 pipeline on all pairwise combinations of replicate populations within a treatment. To calculate the per SNP F_ST_ across time points, we used the Popoolation2 pipeline to compare each population when at passage seven with the ancestor, between passages fourteen and seven, and between passage fourteen and the ancestor. For both analyses we estimated significance using Fisher's Exact Test. We then counted the number of significant loci after Bonferroni corrections. We compared the loci count between populations within each treatment using a chi-square test of goodness-of-fit, and across time points using linear models followed by Tukey multiple-comparison tests to determine pairwise differences.

Analysis of single colony samples

We called variants with the ancestor as the reference using the breseq pipeline with default parameters. We used the output from the breseq gdtools COMPARE command to construct phylogenies with PHYLIP dnapars (*72*). We then used the cophenetic.phylo function of *ape* (*73*) to calculate pairwise distances between the ancestor and each clone. Data were analysed using linear mixed models.

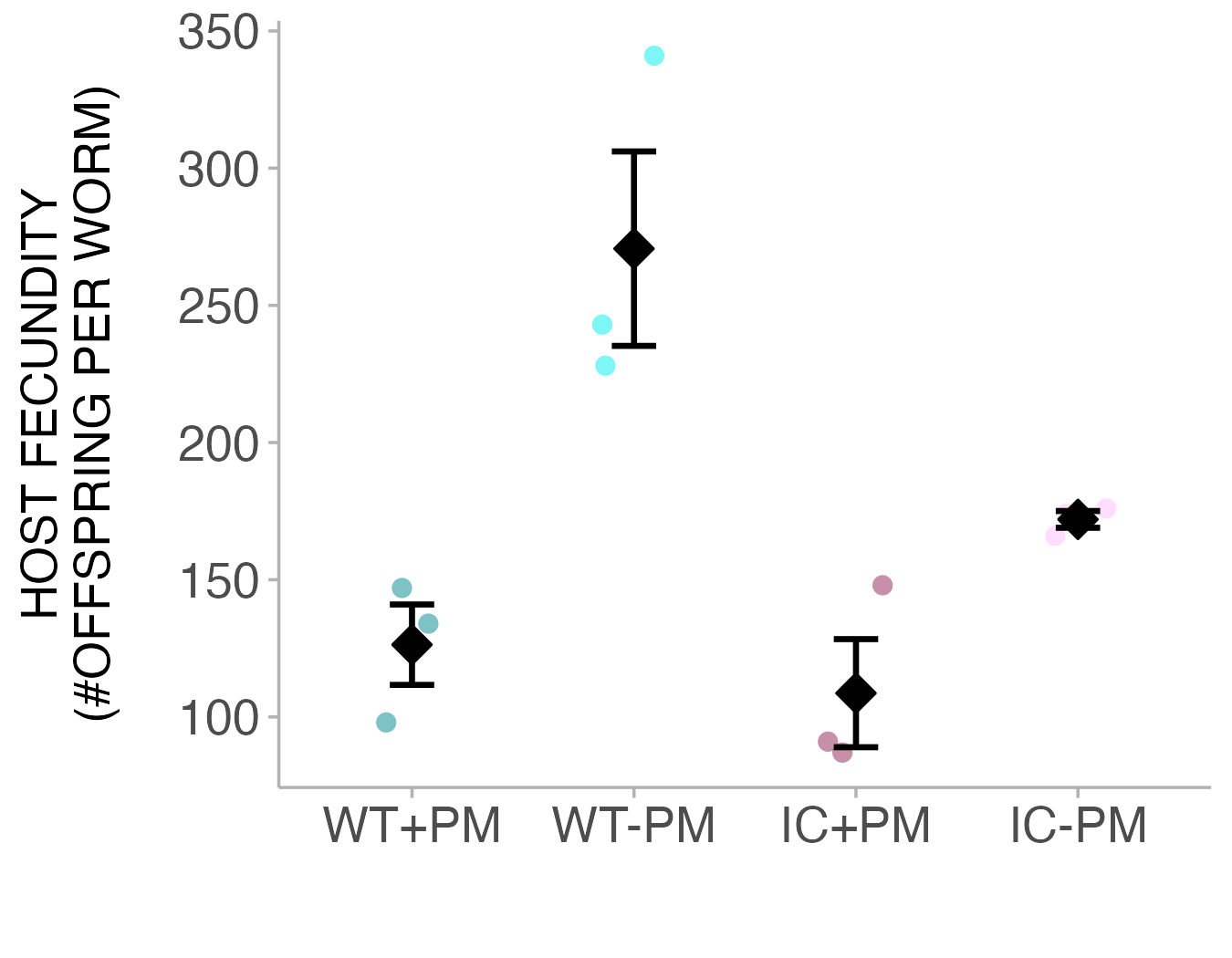

**Figure S1**. Fecundity (mean ± SE) of wild-type and immunocompromised nematodes reared with or without *P. berkeleyensis.* WT = wild-type host, IC = immunocompromised host, PM = protective microbiota.

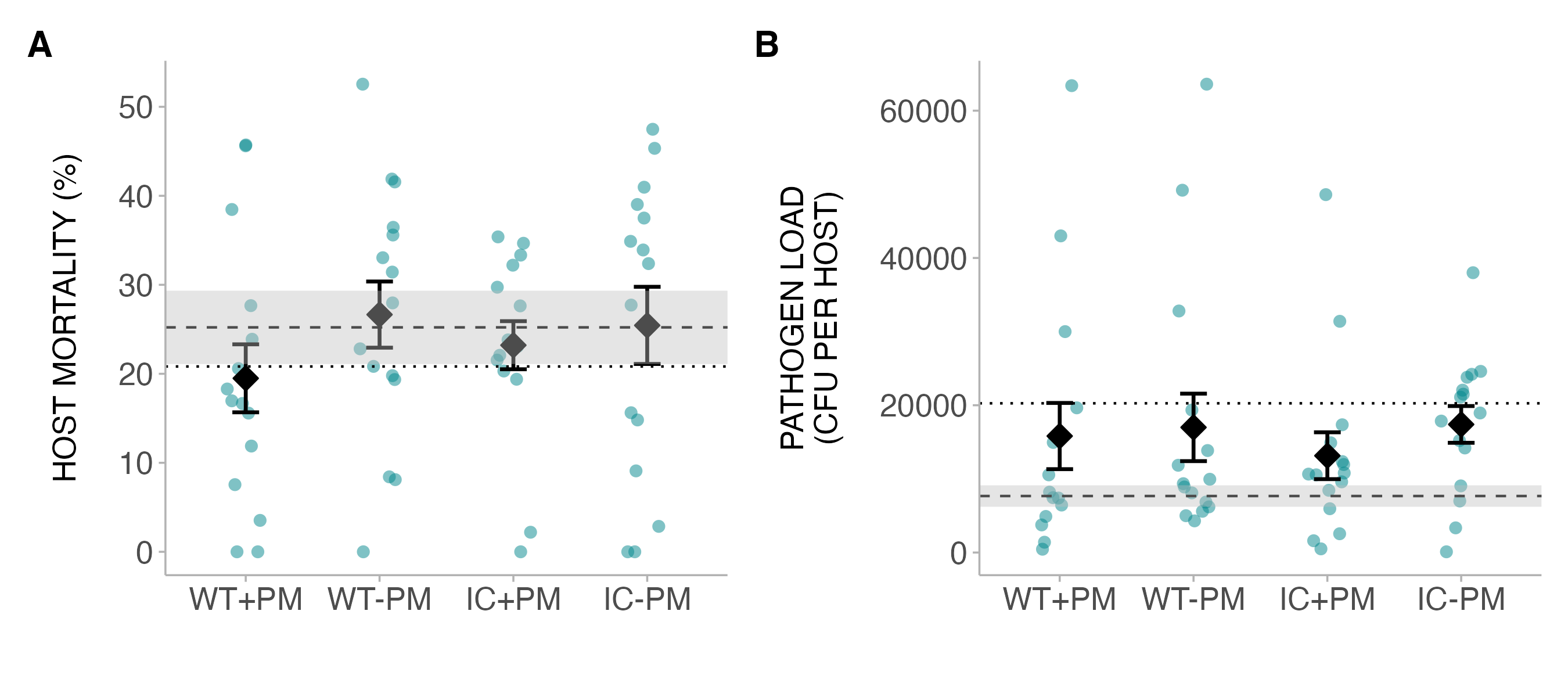

**Figure S2**. (**A**) Mortality (mean ± SE) of wild-type hosts on evolved pathogens (x-axis) after exposure to protective microbiota. (**B**) Load (mean ± SE) of evolved pathogens (x-axis) in wild-type hosts with prior exposure to protective microbiota. Shaded dashed line indicates mean ± SE for hosts infected by no-host control pathogen. Dotted line indicates mean for hosts infected by ancestral pathogen. WT = wild-type host, IC = immunocompromised host, PM = protective microbiota.

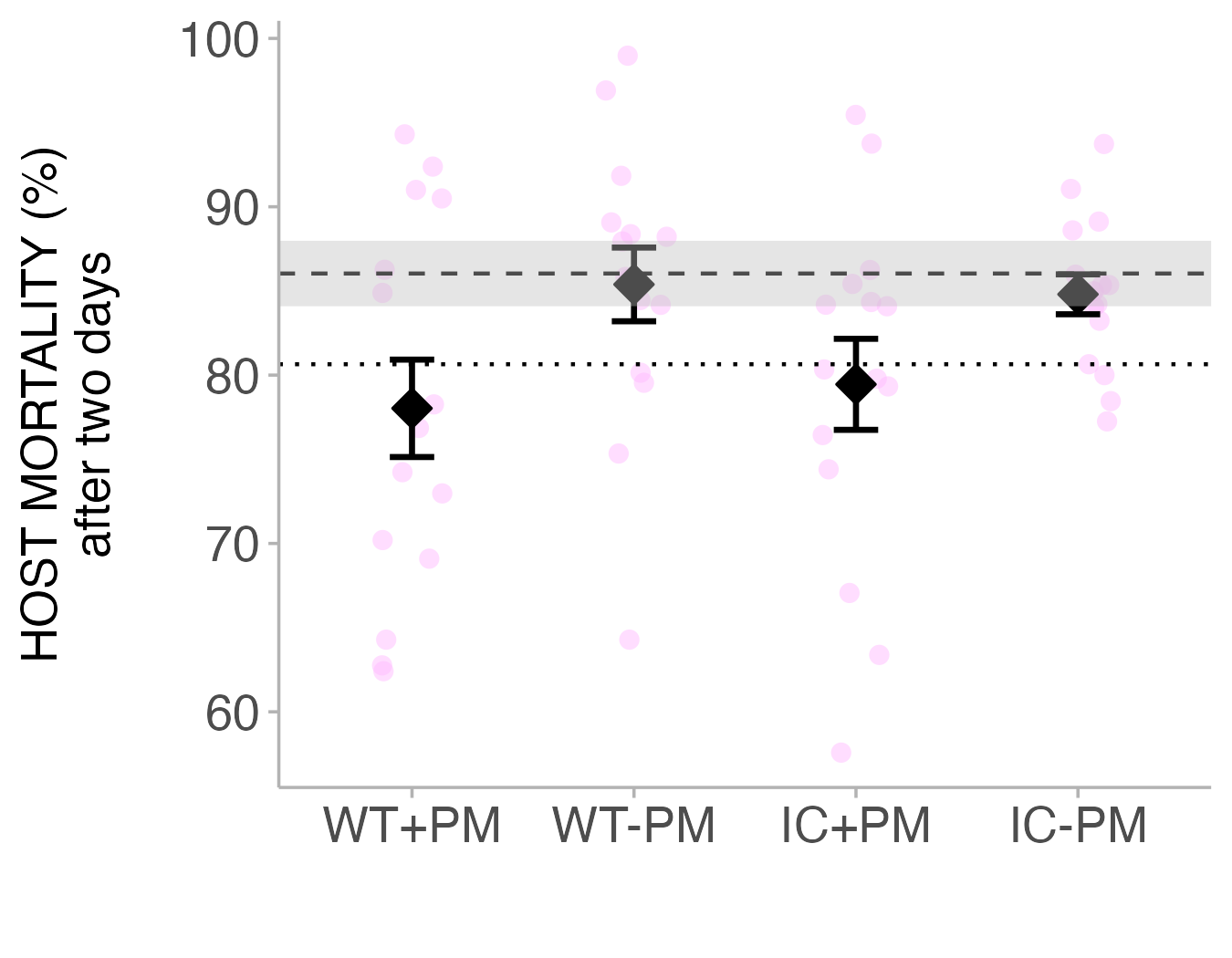

**Figure S3**. Mortality (mean ± SE) of immunocompromised hosts on evolved pathogens (x-axis) without exposure to protective microbiota. Shaded dashed line indicates mean ± SE for hosts infected by no-host control pathogen. Dotted line indicates mean for hosts infected by ancestral pathogen. WT = wild-type host, IC = immunocompromised host, PM = protective microbiota.

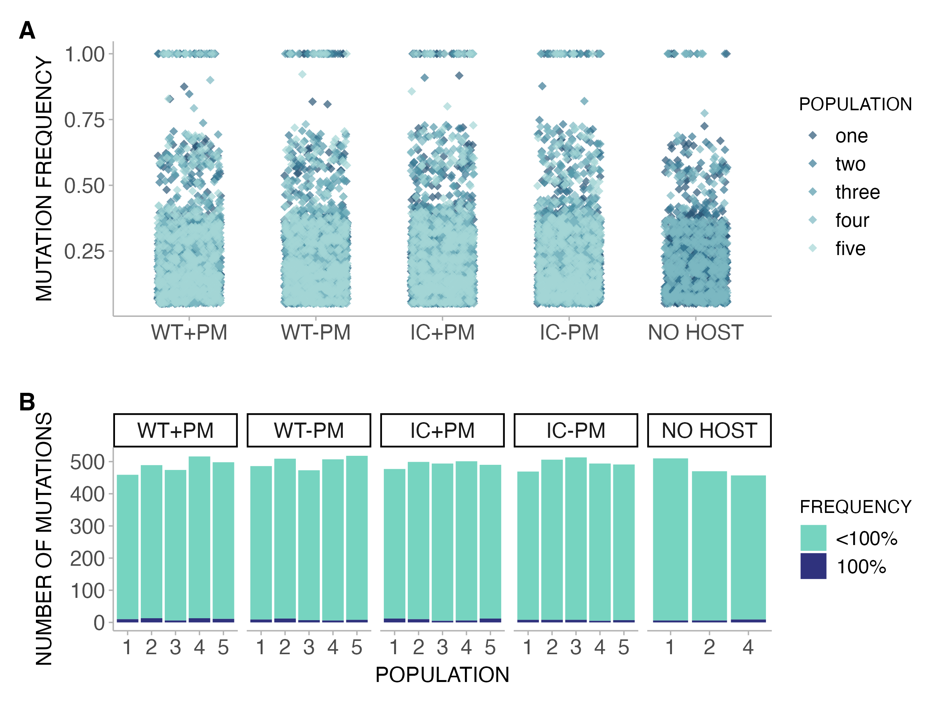

**Figure S4**. (**A**) Distribution of frequencies of mutations in evolved pathogens within each treatment. (**B**) Total number of mutations in evolved pathogens, color-coded by whether mutation is fixed in the population (100%) or not fixed (<100%). WT = wild-type host, IC = immunocompromised host, PM = protective microbiota.

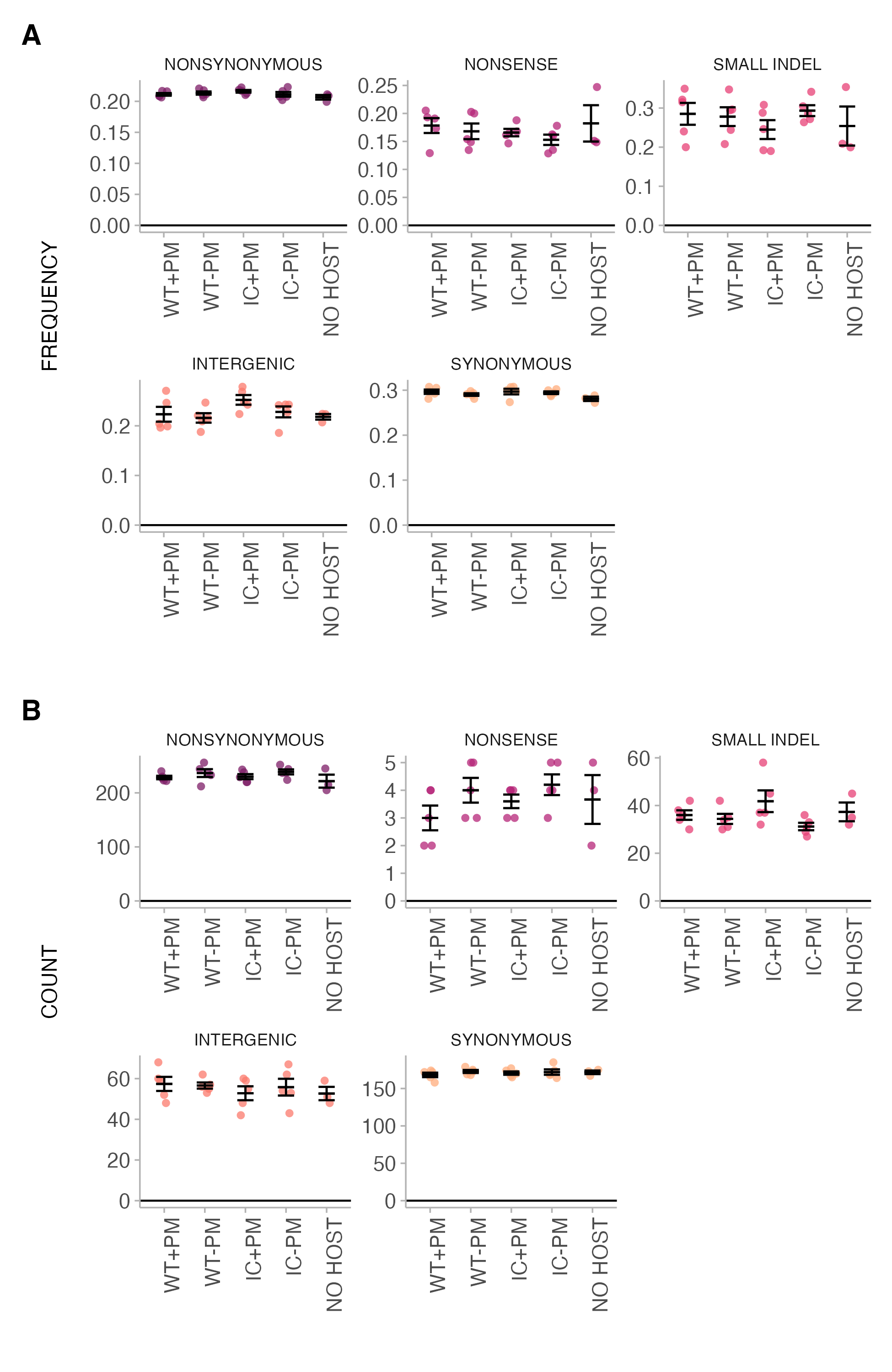

**Figure S5**. (**A**) Frequency and (**B**) count of mutations in evolved pathogen populations grouped by mutation category. All errors bars are mean ± SE. WT = wild-type host, IC = immunocompromised host, PM = protective microbiota.

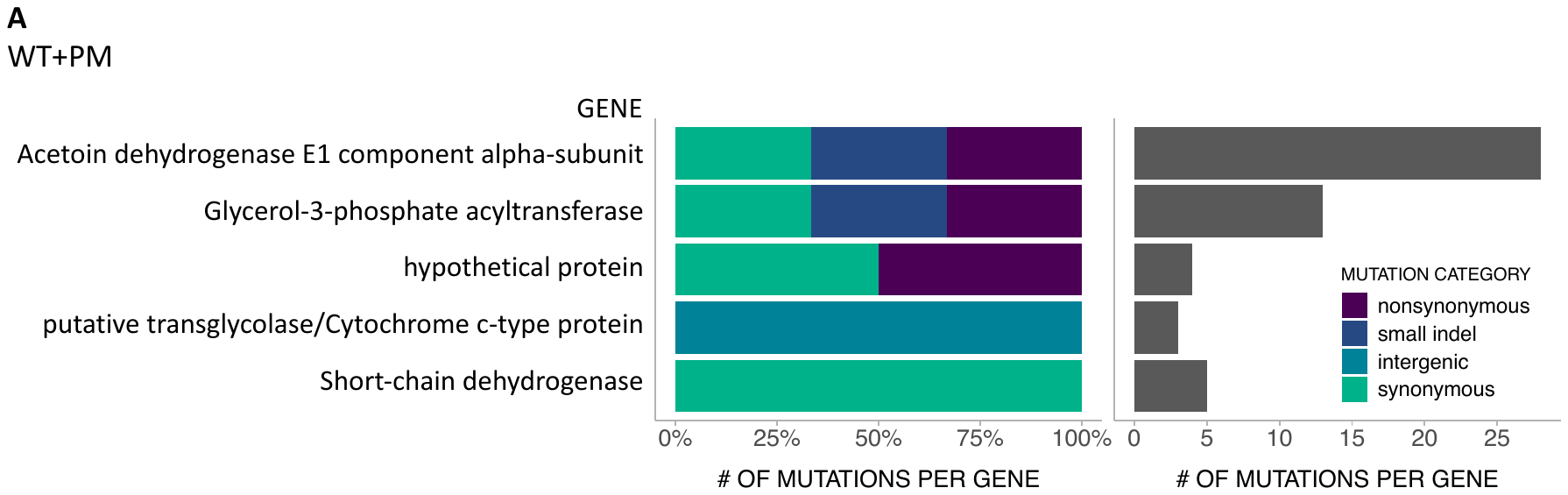

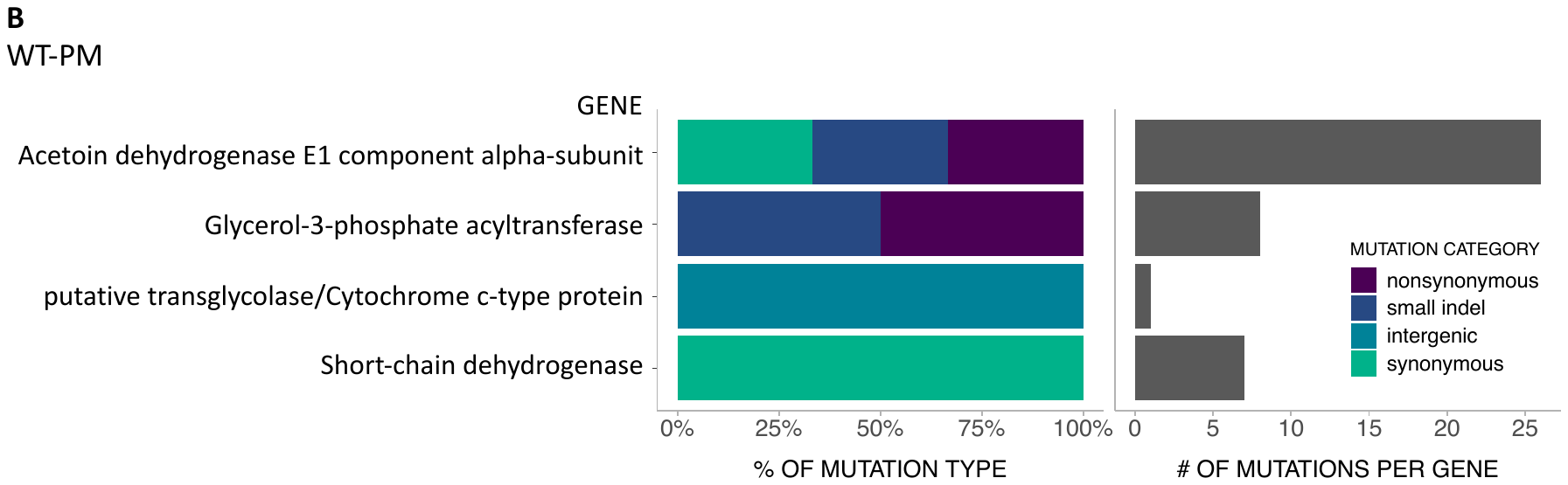

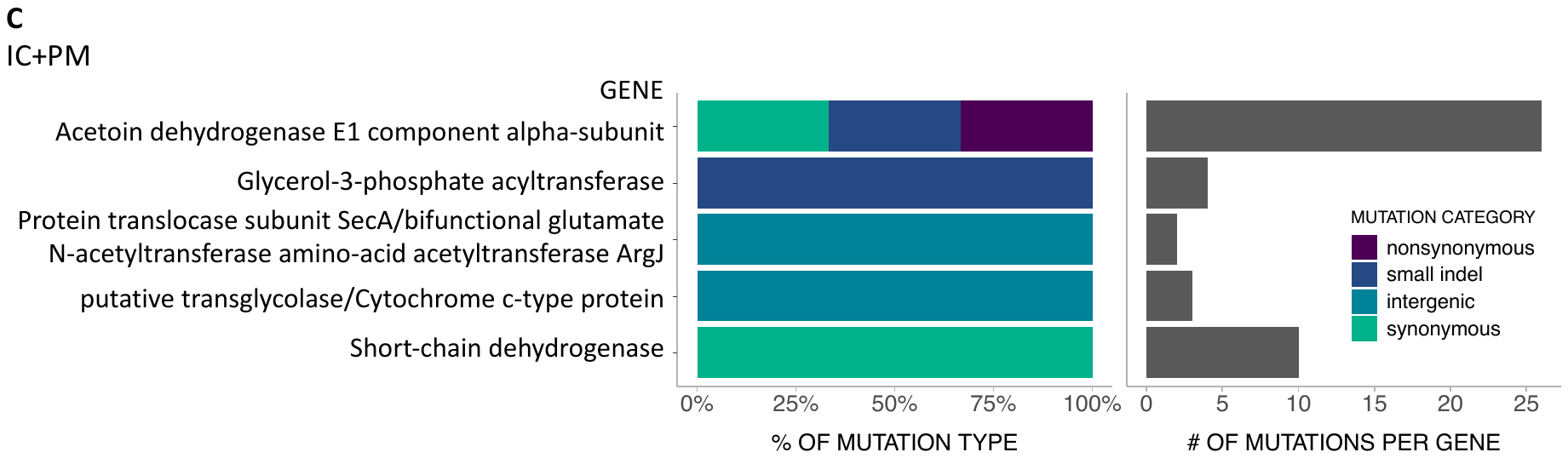

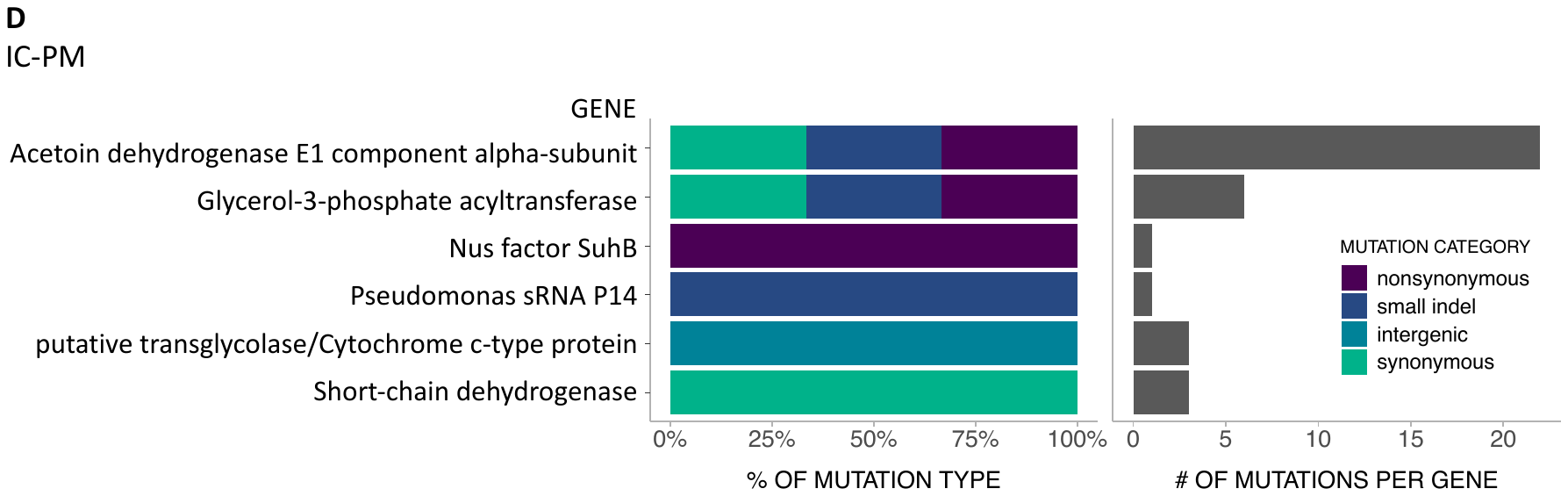

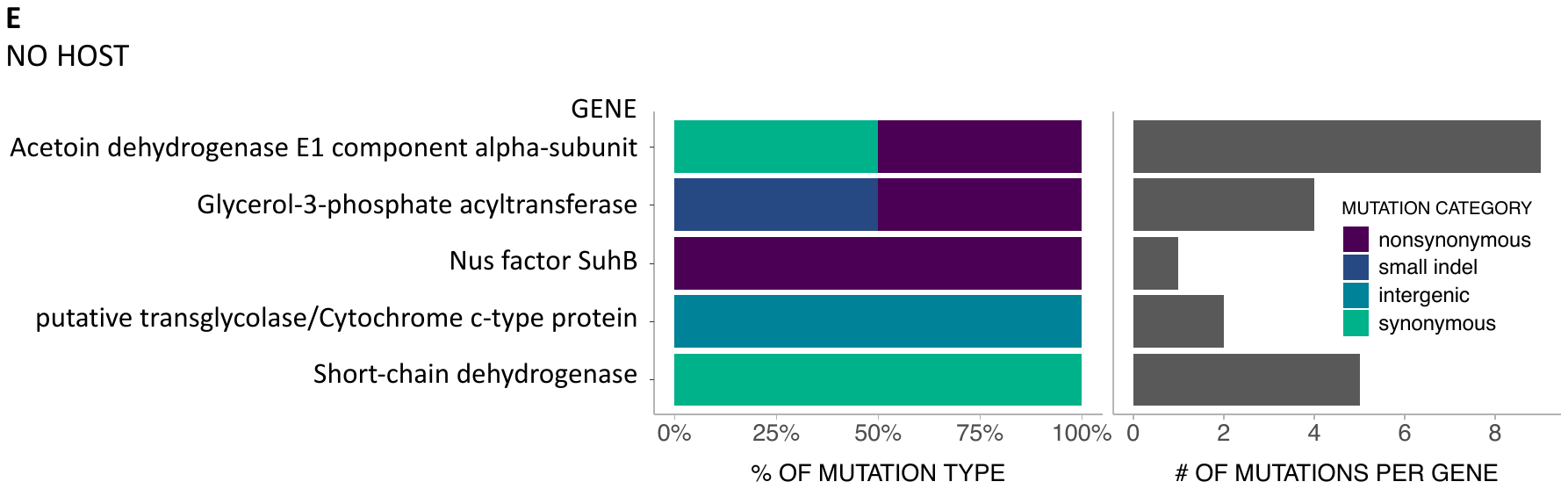

**Figure S6**. (**A-E**) Genes where mutations have fixed in at least one population within each treatment. “Percent of mutation type” indicates the proportion of the “number of mutations per gene” belonging to respective mutation category. WT = wild-type host, IC = immunocompromised host, PM = protective microbiota.

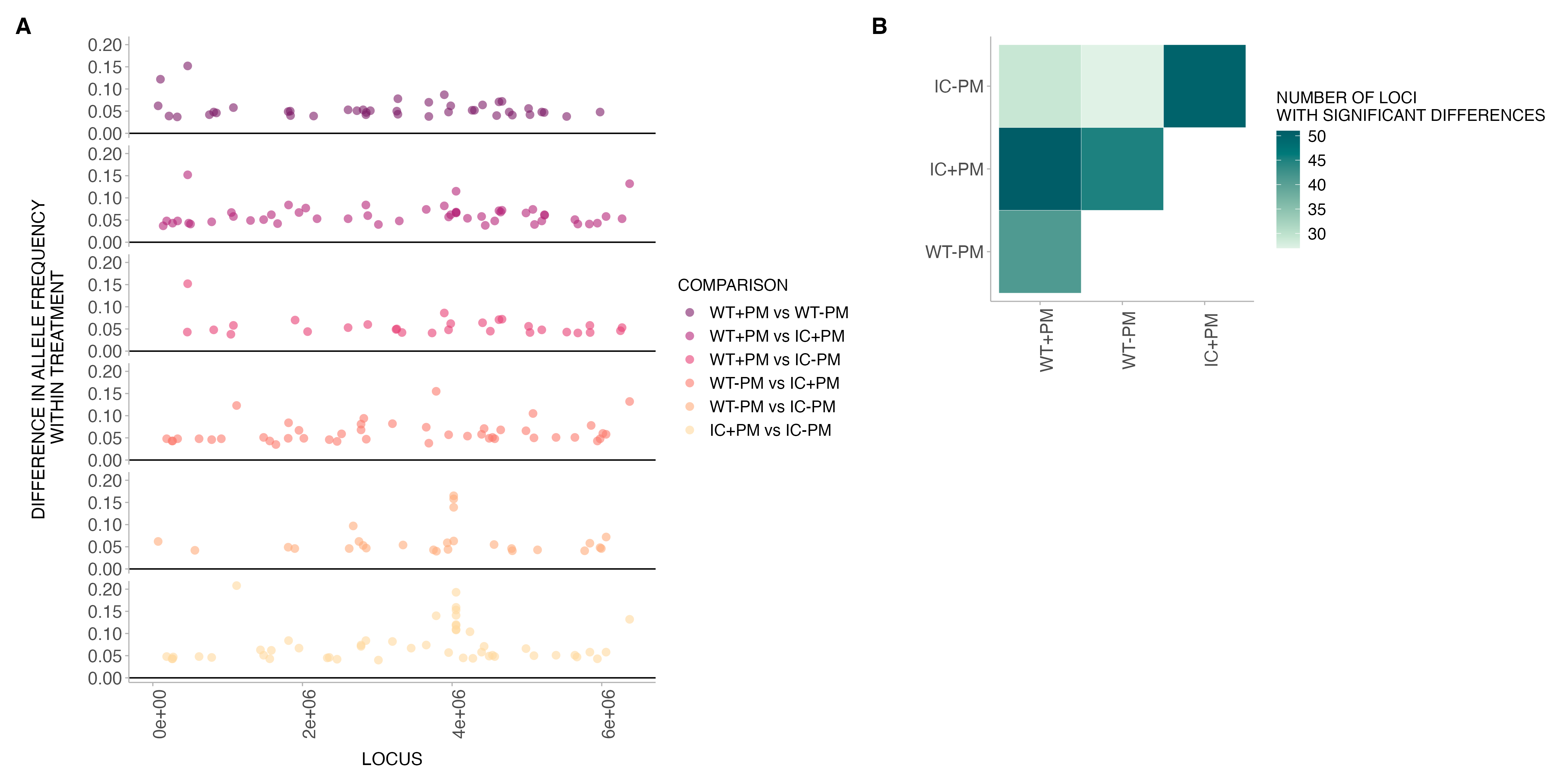

**Figure S7**. (**A**) Relative position of mutations where there were significant differences in allele frequency between treatments (**B**) Total number of loci in (A) with significant differences in allele frequency between treatments. WT = wild-type host, IC = immunocompromised host, PM = protective microbiota.

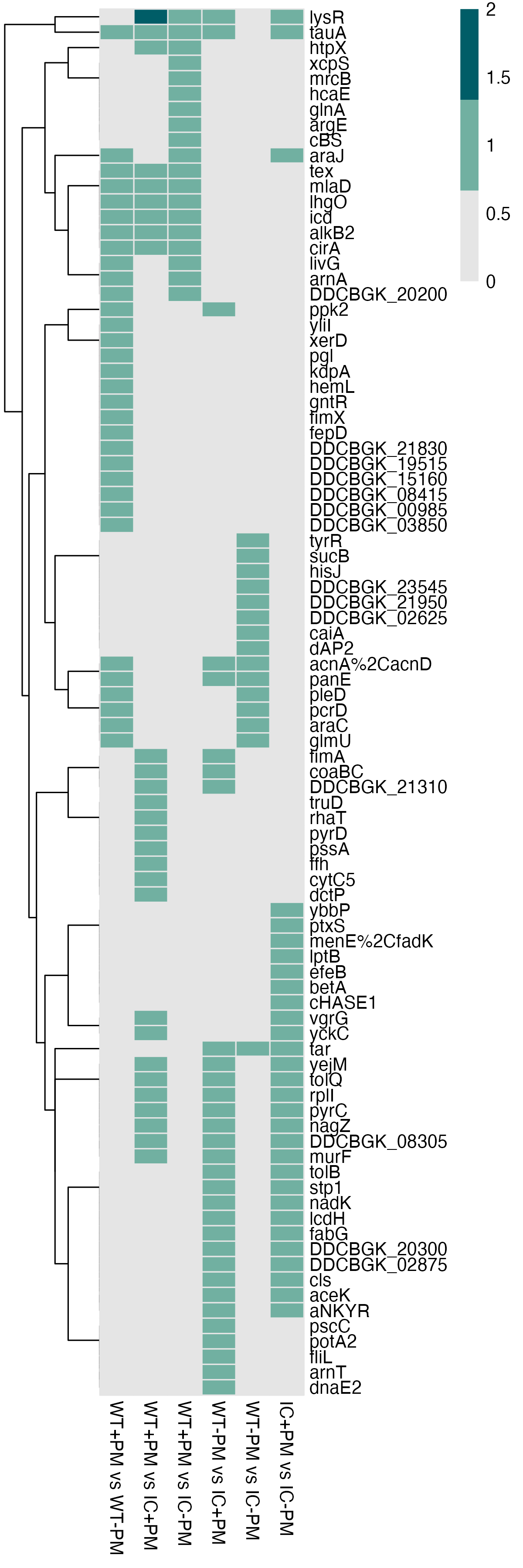

**Figure S8** (**A**). Heatmap of genes with significant differences in allele frequency between treatments from Figure S7. WT = wild-type host, IC = immunocompromised host, PM = protective microbiota.

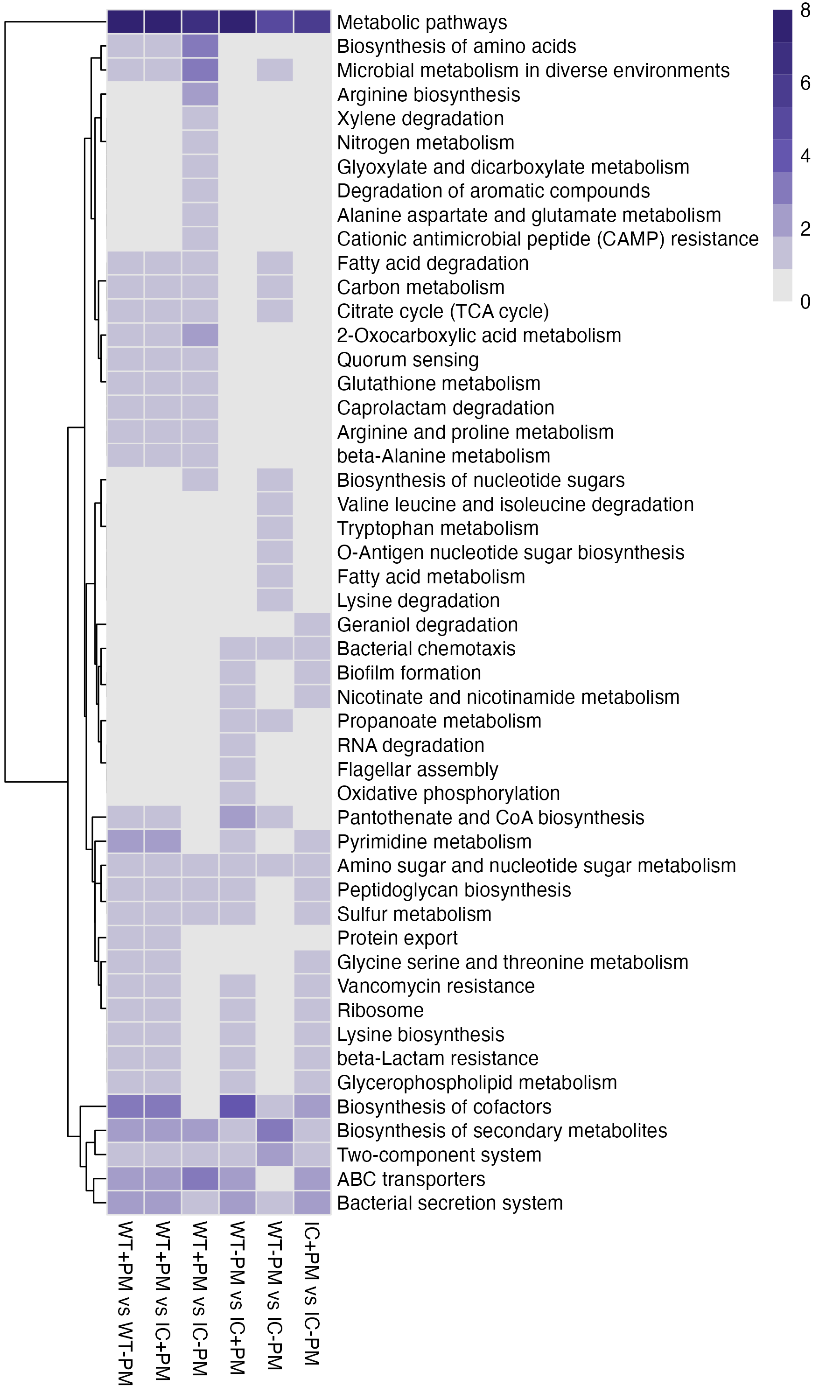

**Figure S8** (**B**). Heatmap of genes from (A) mapped to KEGG pathways. Colour gradient indicates number of genes in each KEGG term. WT = wild-type host, IC = immunocompromised host, PM = protective microbiota.

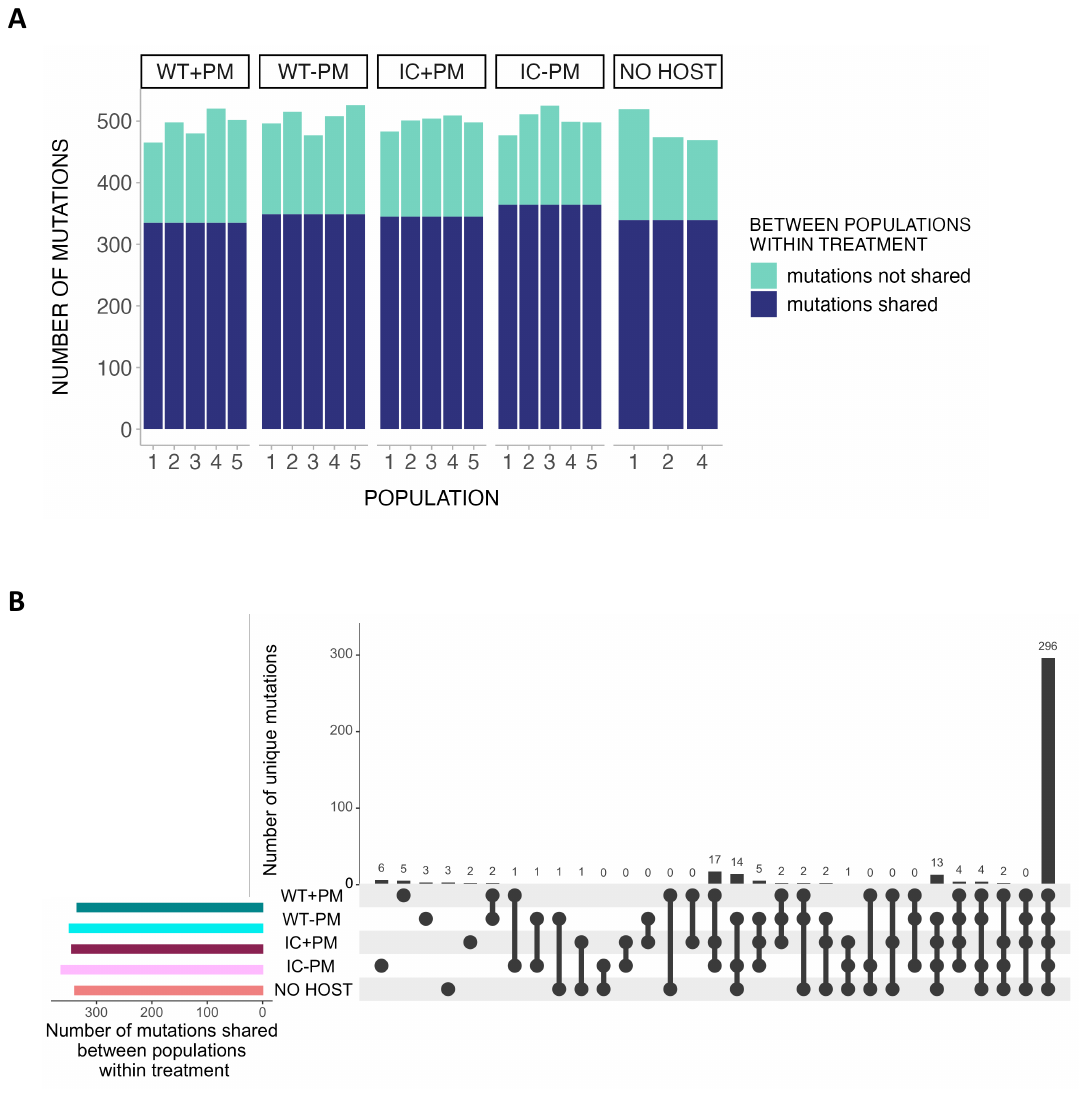

**Figure S9**. (**A**) Number of mutations in common or not in common across all replicate populations within each treatment (**B**) Out of all the mutations in common within each treatment (dark blue portion of ((A)), the number of mutations unique to respective treatment or treatments. For example, there are 6 mutations unique to the WT+PM treatment not found in other treatments, and 0 mutations in common between IC-PM and no host that are unique to these two treatments. WT = wild-type host, IC = immunocompromised host, PM = protective microbiota.

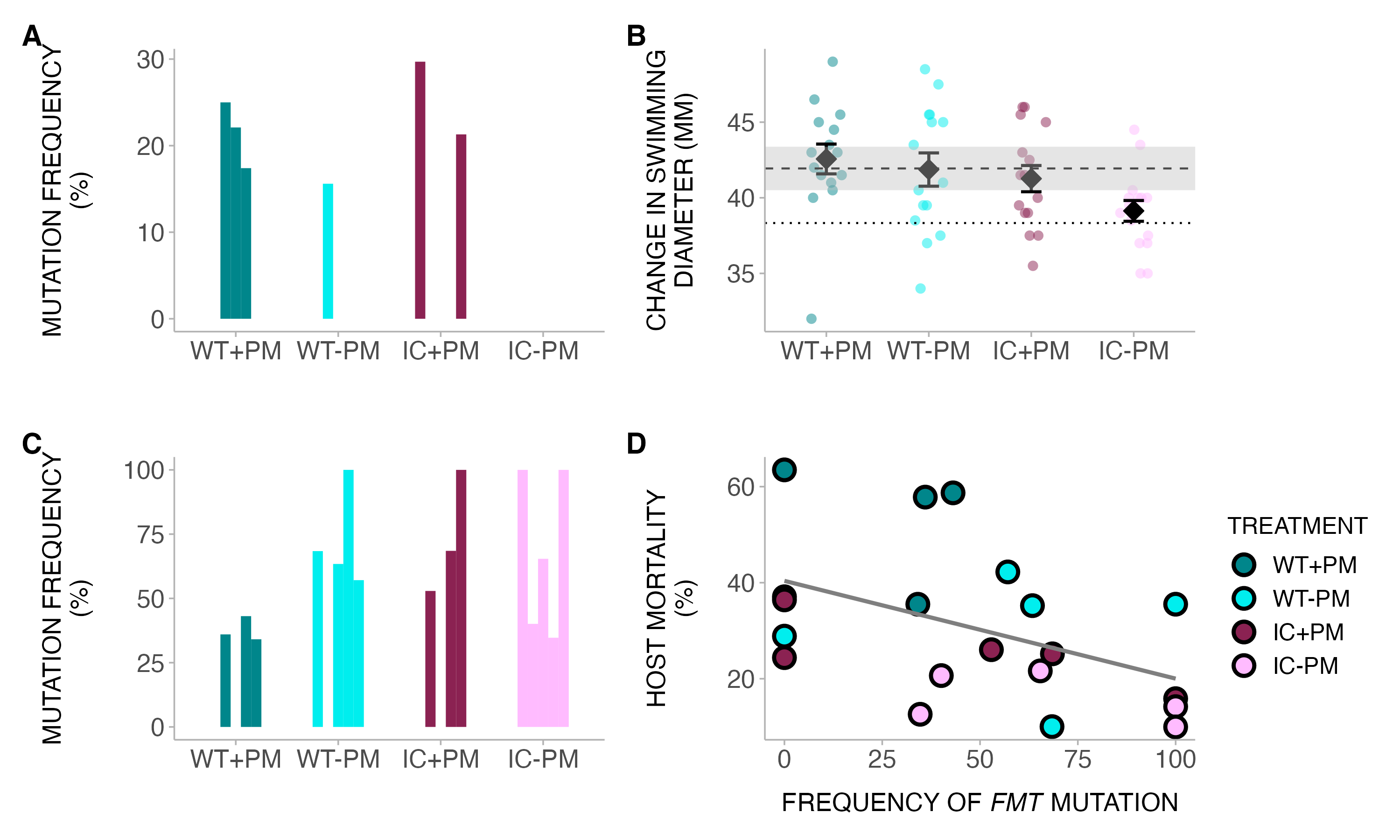

**Figure S10**. (**A**) Frequency of *flge*/*flgf* mutation in respective treatment. Each bar represents one evolved pathogen population (**B**) Swimming motility (mean ± SE) of evolved pathogens. Shaded dashed line indicates mean ± SE of no host treatment. Dotted line indicates mean of ancestral pathogen. (**C**) Frequency of *fmt* mutation in respective treatment. Each bar represents one evolved pathogen population (**D**) Correlation between host mortality and *flge*/*flgf* mutation. WT = wild-type host, IC = immunocompromised host, PM = protective microbiota.

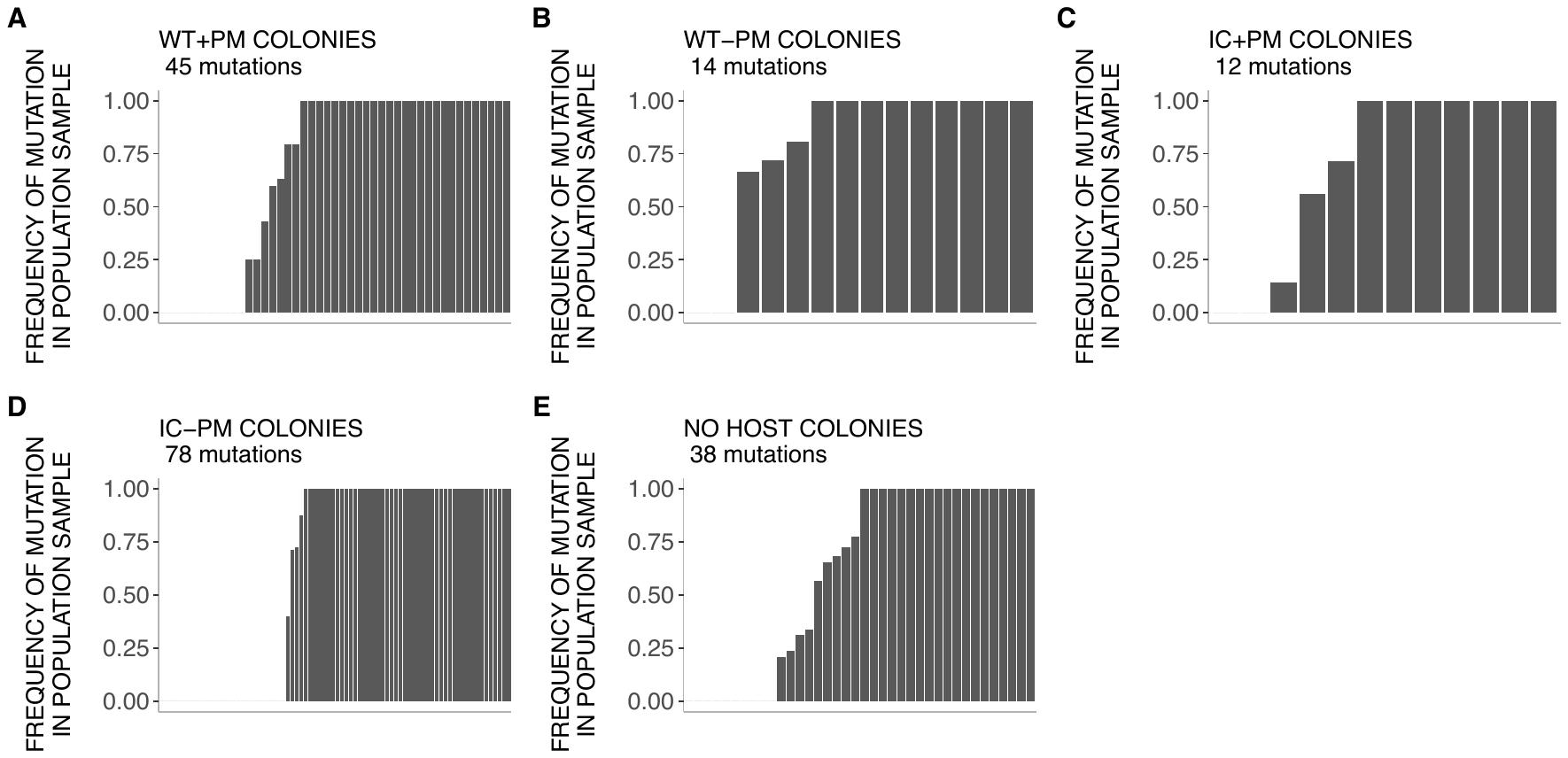

**Figure S11**. (**A-E**) Frequency of mutations in pooled samples shared with single colony samples. The total number of mutations across all clones sampled are indicated under the treatment name. We sampled more clones from WT+PM and IC-PM treatments (3 per population x 5 populations) than the other treatments (1 per population x 5 populations for WT-PM and IC+PM, and 1 per population x 3 populations for no host treatment). WT = wild-type host, IC = immunocompromised host, PM = protective microbiota.

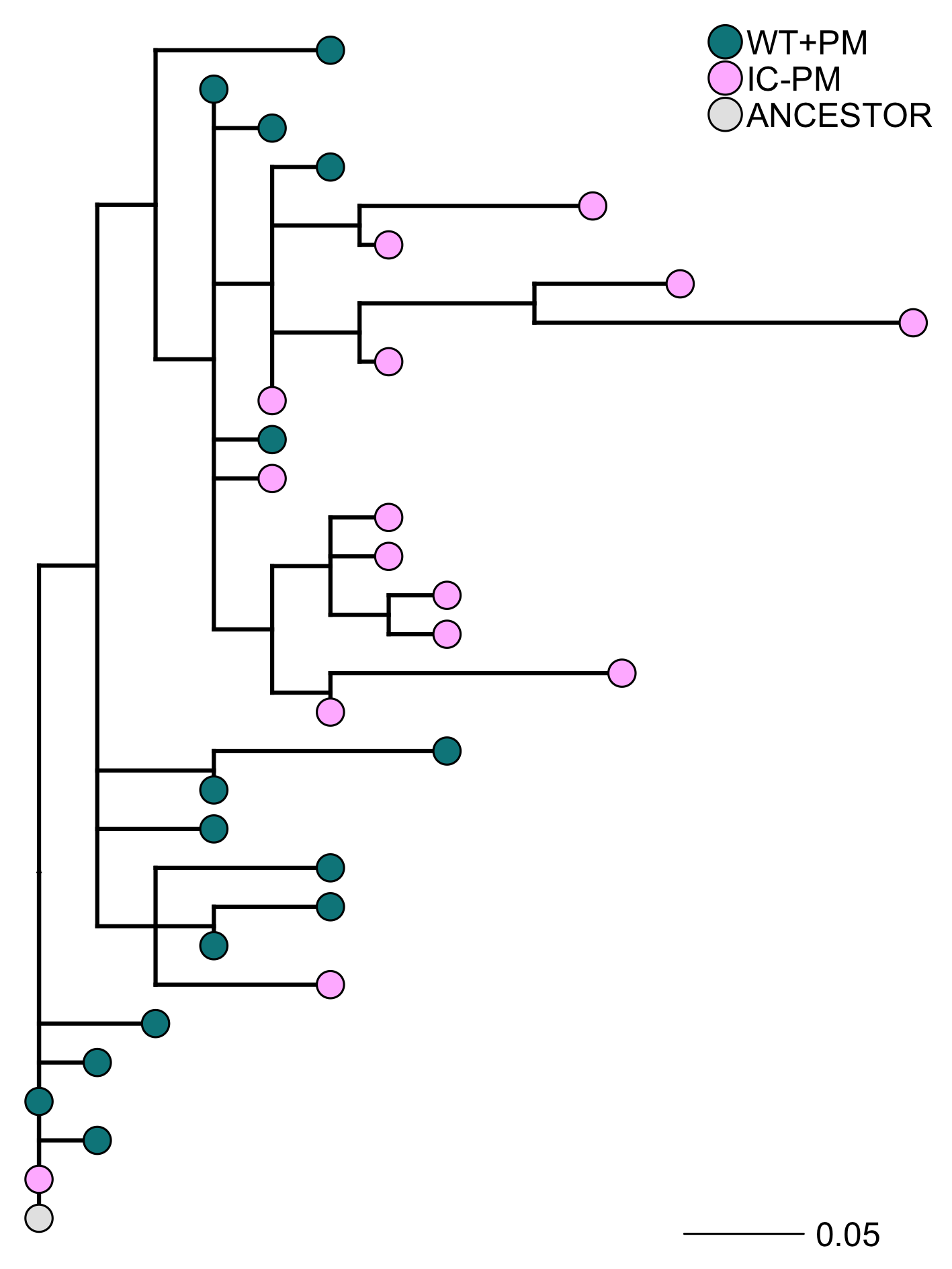

**Figure S12**. Maximum parsimony phylogeny of colonies sampled from WT+PM and IC-PM treatments. WT = wild-type host, IC = immunocompromised host, PM = protective microbiota.

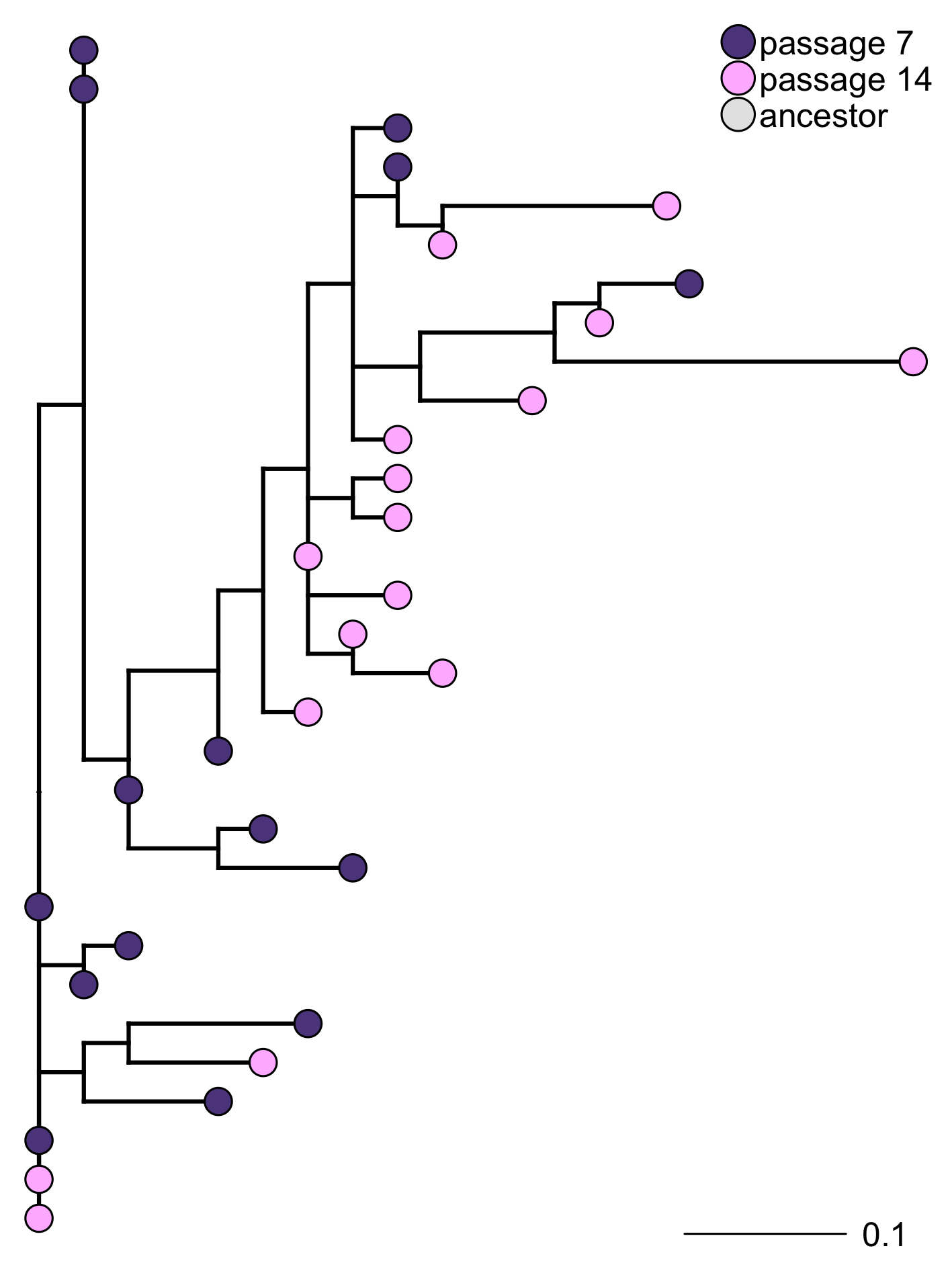

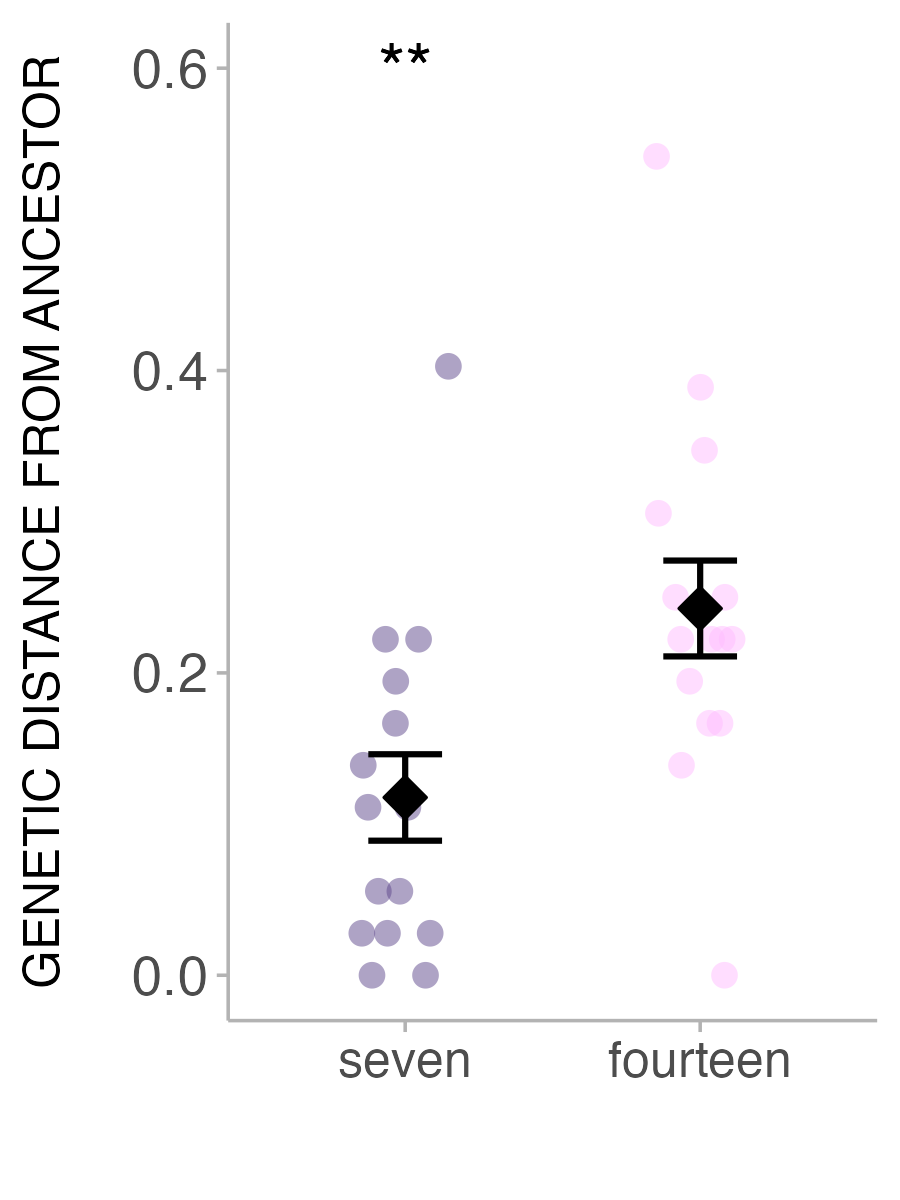

**Figure S13**. Maximum parsimony phylogeny of colonies sampled from passages 7 and 14 of immunocompromised hosts without protective microbiota treatment. (inset) Genetic distance from the ancestor (mean ± SE) for colonies isolated from passages 7 and 14. Error bars indicate standard errors. **P > 0.01

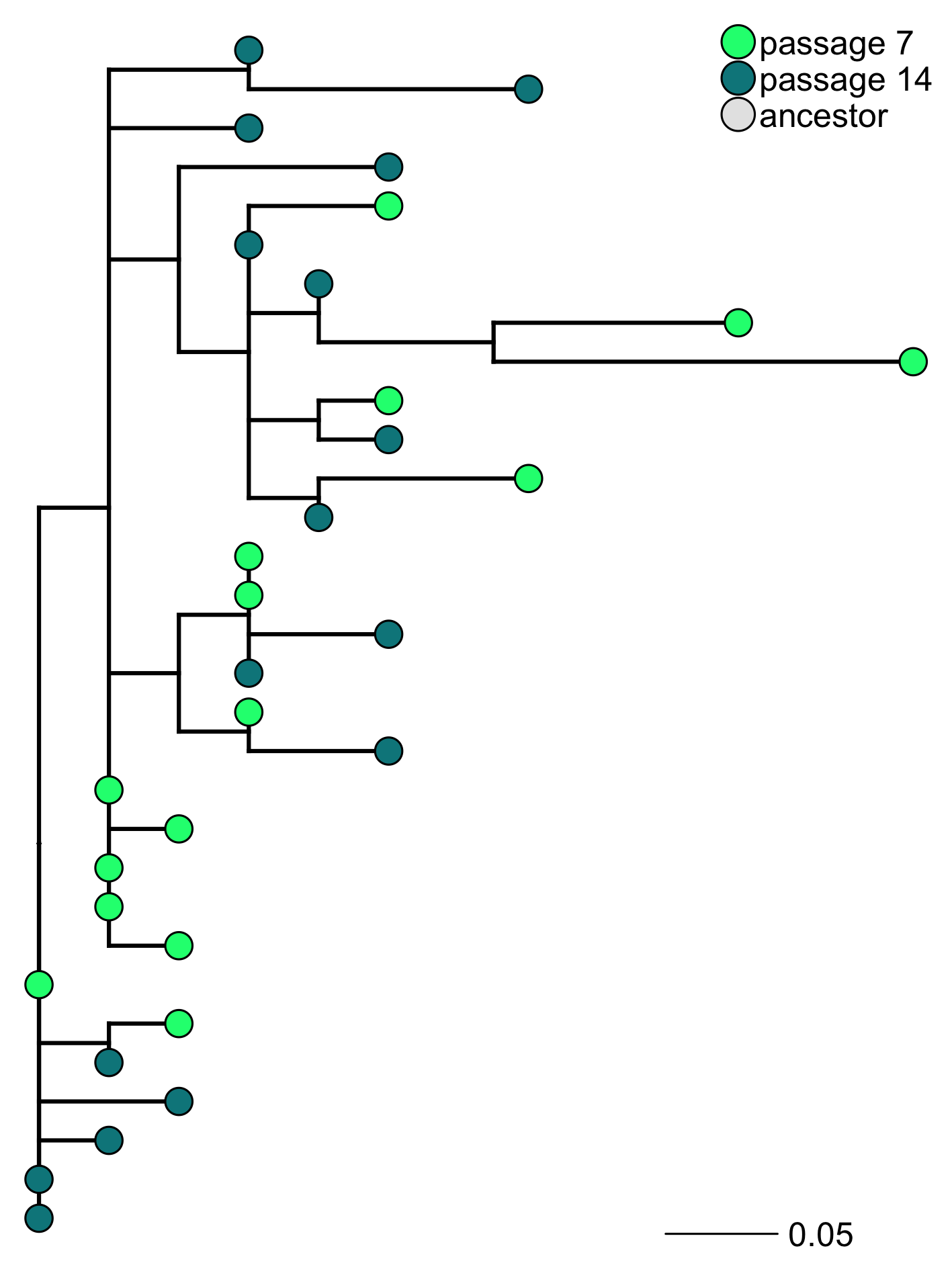

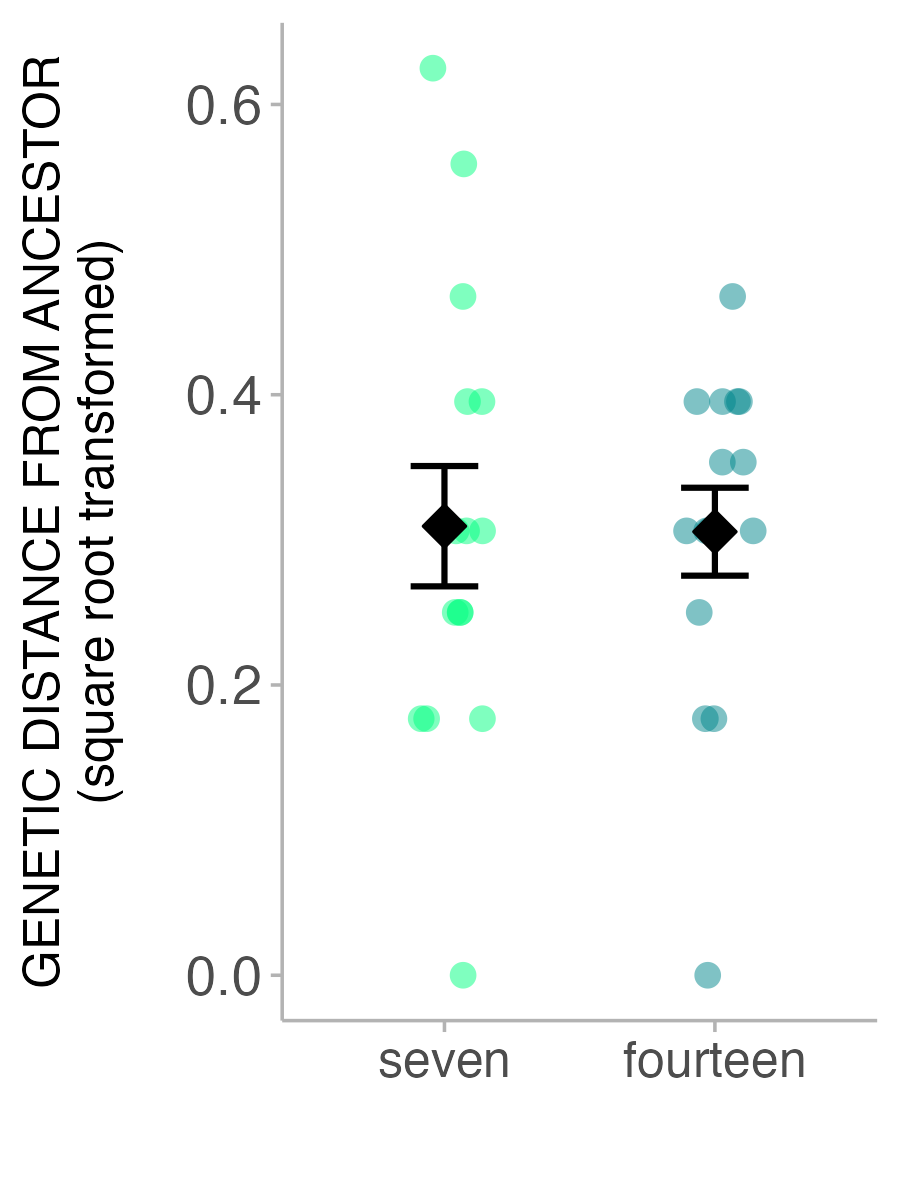

**Figure S14**. Maximum parsimony phylogeny of colonies sampled from passages 7 and 14 of wild-type hosts with protective microbiota treatment. (inset) Genetic distance from the ancestor (mean ± SE) for colonies isolated from passages 7 and 14. Error bars indicate standard errors.

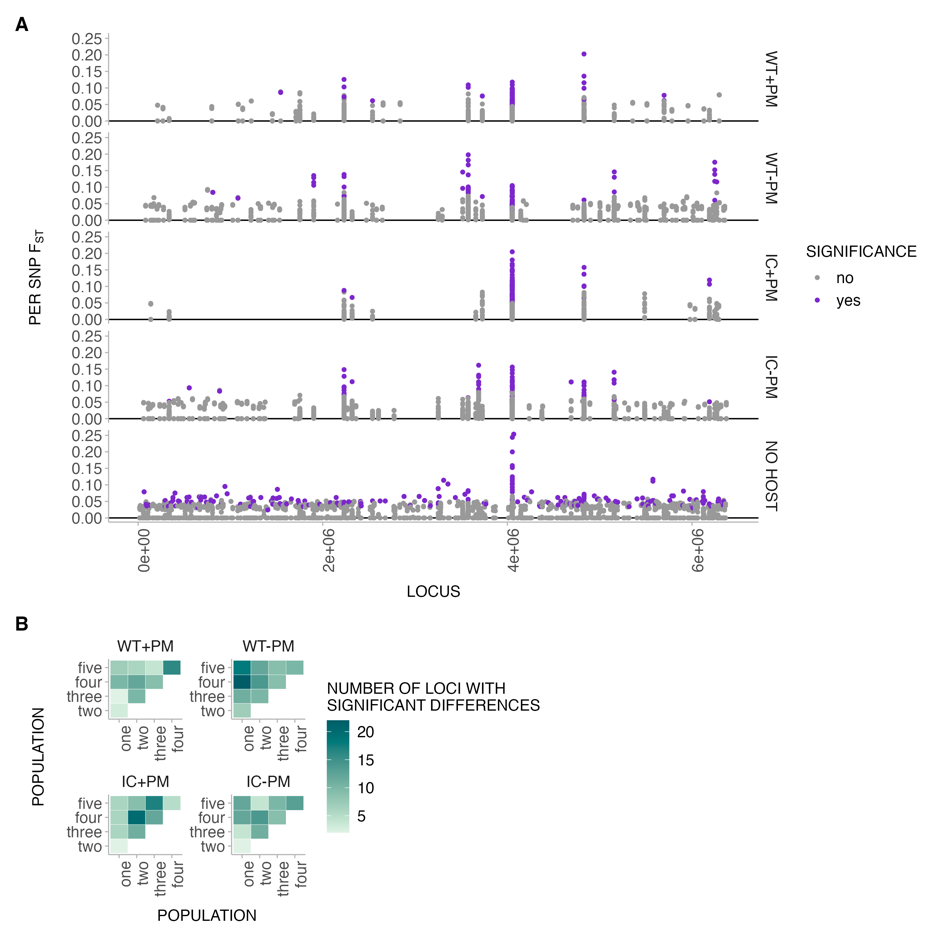

**Figure S15.** (**A**) Per SNP F_ST_ across pathogen genome for pairwise comparisons of populations within each treatment, where purple dots indicate significant F_ST_ after Bonferroni correction (**B**) Total number of loci in (A) with significant F_ST_ after Bonferroni correction. WT = wild-type host, IC = immunocompromised host, PM = protective microbiota.

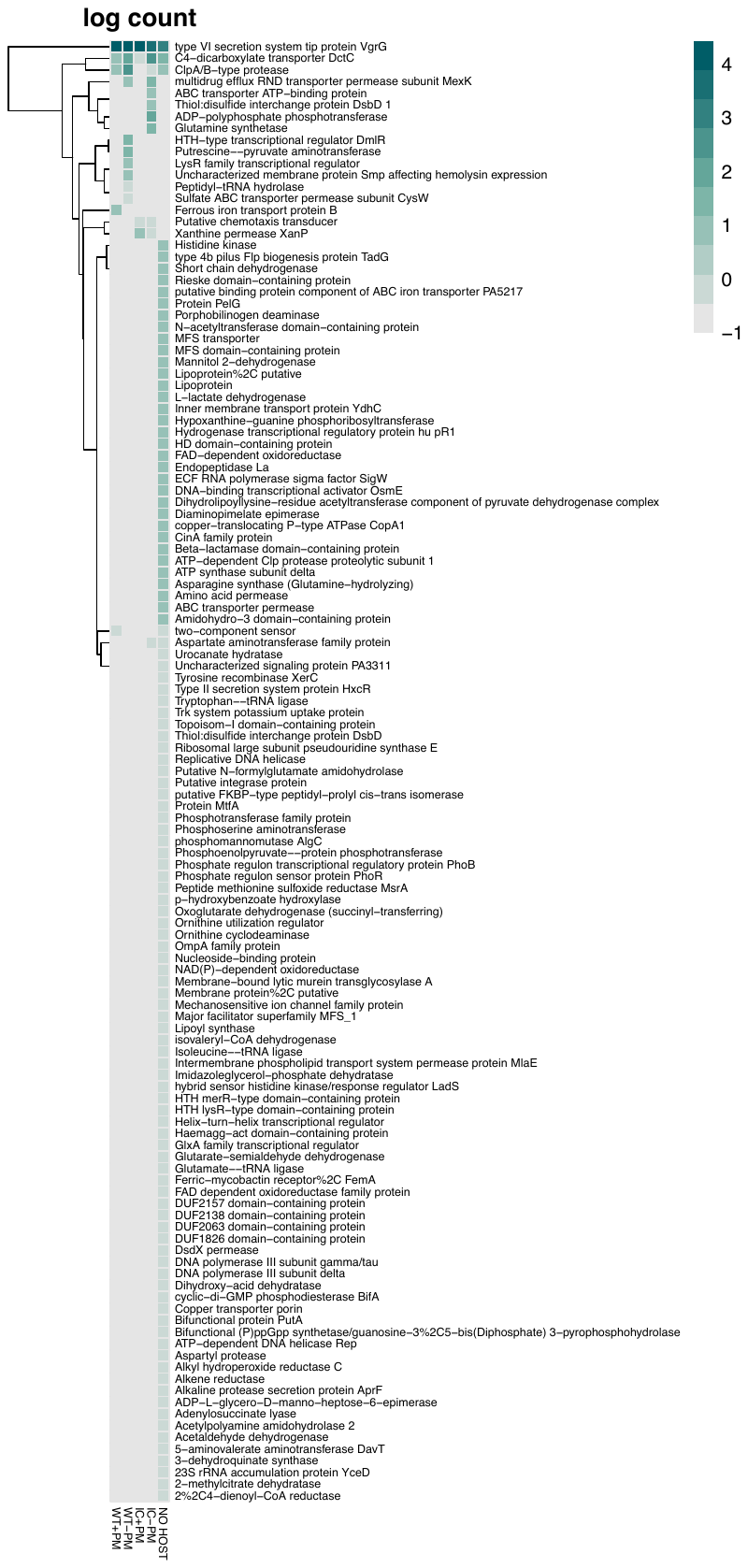

**Figure S16A**. Heatmap of genes with significant differences in per SNP F_ST_ from Figure S15. Colours indicate the natural log of the gene count. WT = wild-type host, IC = immunocompromised host, PM = protective microbiota.

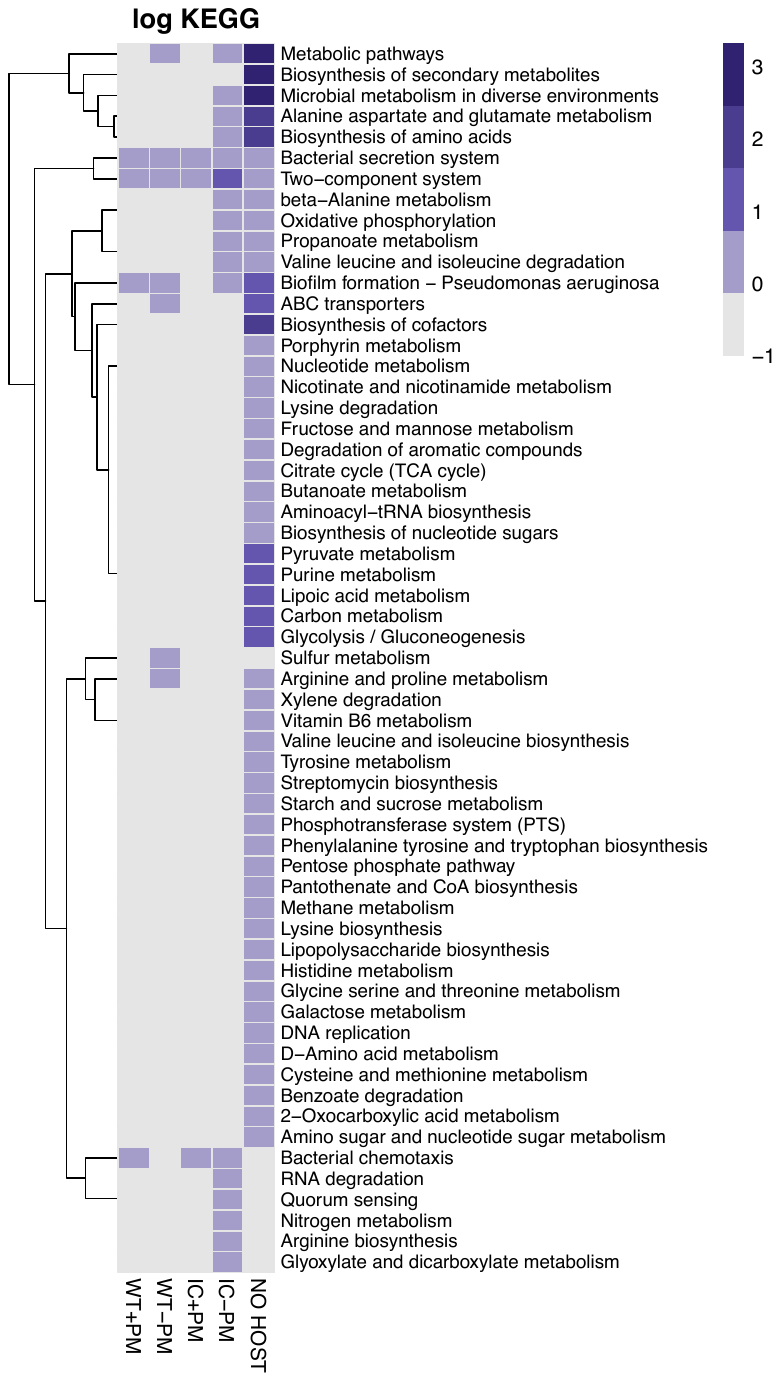

**Figure S16** (**B**). Heatmap of genes from (A) mapped to KEGG pathways. Colour gradient indicates the natural log of the number of genes in each KEGG term. WT = wild-type host, IC = immunocompromised host, PM = protective microbiota.

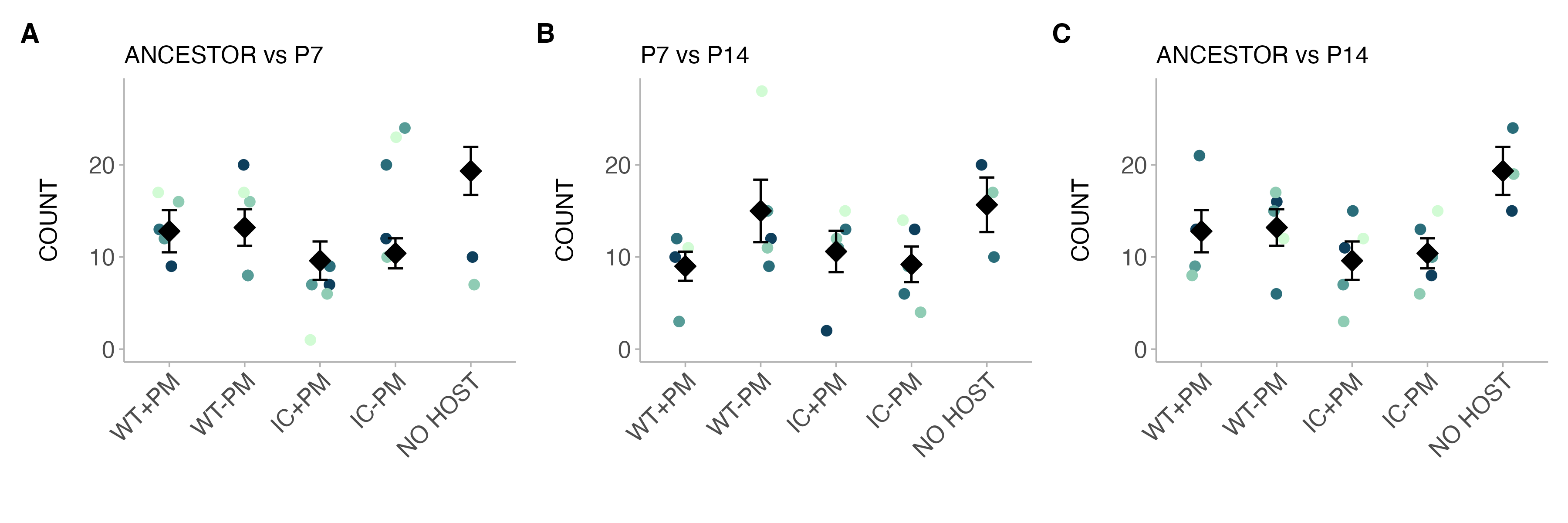

**Figure S17**. Count (mean ± SE) of significant per SNP F_ST_ between (**A**) the ancestor and passage 7, (**B**) passage 7 and passage 14, and (**C**) the ancestor and passage 14.
